# Shelterin components modulate nucleic acids condensation and phase separation in the context of telomeric DNA

**DOI:** 10.1101/2021.04.30.442189

**Authors:** Andrea Soranno, J. Jeremías Incicco, Paolo De Bona, Eric J. Tomko, Eric A. Galburt, Alex S. Holehouse, Roberto Galletto

## Abstract

Telomeres are nucleoprotein complexes that protect the ends of chromosomes and are essential for chromosome stability in Eukaryotes. In cells, individual telomeres form distinct globules of finite size that appear to be smaller than expected for bare DNA. Moreover, upon changes in their protein composition, telomeres can cluster to form telomere-induced-foci (TIFs) or co-localize with promyelocytic leukemia (PML) nuclear bodies. The physical basis for collapse of individual telomeres and coalescence of multiple ones remains unclear, as does the relationship between these two phenomena. By combining single-molecule measurements, optical microscopy, turbidity assays, and simulations, we show that the telomere scaffolding protein TRF2 can condense individual DNA chains and drives coalescence of multiple DNA molecules, leading to phase separation and the formation of liquid-like droplets. Addition of the TRF2 binding protein hRap1 modulates phase boundaries and tunes the specificity of solution demixing while simultaneously altering the degree of DNA compaction. Our results suggest that the condensation of single telomeres and formation of biomolecular condensates containing multiple telomeres are two different outcomes driven by the same set of molecular interactions. Moreover, binding partners, such as other telomere components, can alter those interactions to promote single-chain DNA compaction over multiple-chain phase separation.

## Introduction

Biomolecular condensates are non-stochiometric assemblies of biomacromolecules that underlie a variety of cellular processes including the stress response, ribosomal assembly, germ-granule formation and chromatin organization (1-8). These assemblies are commonly enriched in multivalent DNA/RNA binding proteins and nucleic acids, with multivalency being a critical feature that facilitates formation of a set of reversible inter-molecular interactions (9-13). As a result of this multivalency, many of the proteins that localize in condensates have been shown to undergo phase separation *in vitro* and *in vivo*, lending support to a model in which intracellular phase separation plays a key role in cellular organization and compartmentalization (14-17).

The emergent property of biomolecular condensates parallels previously observed characteristics of ligand-induced condensation of nucleic acids, in which multivalent interactions provided by DNA/RNA binding proteins can promote either the compaction of single nucleic acid chains or the condensation of multiple ones. Indeed, as proposed by Post and Zimm (18), from a polymer physics standpoint, the condensation of a single nucleic acid chain by a ligand and the phase separation of mixtures of nucleic acids and the same ligands can be rationalized as two distinct outcomes of the same set of interactions. Intriguingly, Post and Zimm (18) proposed the investigation of phase separation as a method to test molecular interactions within single nucleic acids, since at the time access to the experimental study of single molecule conformations and dynamics was not available. Advancements in single-molecule methods and their application have paved the way to a direct investigation of protein-mediated nucleic acid condensation (19-24). While the relative concentration regimes and molecular architecture of the components play key roles, the physical basis that determines whether protein-nucleic acids interactions lead to intra-molecular loops and chain compaction or to inter-chain interactions and solution demixing remains to be fully elucidated. An important implication of this duality is that, under appropriate conditions, many proteins that can induce the condensation of single nucleic-acid chains will instead promote phase separation. Though rooted into a similar theoretical framework (18, 25), this is a distinct phenomenon from the compaction and phase separation observed for a single protein (26-28), since it entails the interaction between a nucleic acid (often significantly long) and its interaction with ligands. Identifying how these two possible outcomes are encoded in the architecture of proteins and their interaction with specific and unspecific nucleic acids is fundamental to our understanding of how the genetic material may be organized and regulated *in vivo*. To address this question directly we focused on the telomeric protein TRF2 and sought to explore the symmetry between nucleic acid compaction and phase separation.

Telomeres are nucleoprotein complexes that protect the ends of linear chromosomes. Human telomeres are formed by the six shelterin subunits TRF1, TRF2, Rap1, TIN2, TPP2 and POT1 that co-assemble at the end of chromosomes (29) (Fig. 1A), with the four-protein core complex TRF1-TIN2-TRF2-Rap1 forming the major scaffold bound to the terminal double-stranded DNA containing T_2_AG_3_ repeats. Homodimers of TRF1 and TRF2 bind specifically to the telomeric repeats (30-32), while Rap1 interacts with TRF2 with high affinity (33, 34), and TIN2 bridges TRF1 and TRF2 (35, 36). Furthermore, TIN2 interacts with TPP1-POT1 bound to the G-rich 3’ single-stranded DNA overhang, providing a physical link with the core complex formed on the double-stranded DNA telomeric region and thus stabilizing a T-loop structure proposed to regulate telomere accessibility (37-39).

**Figure 1.**
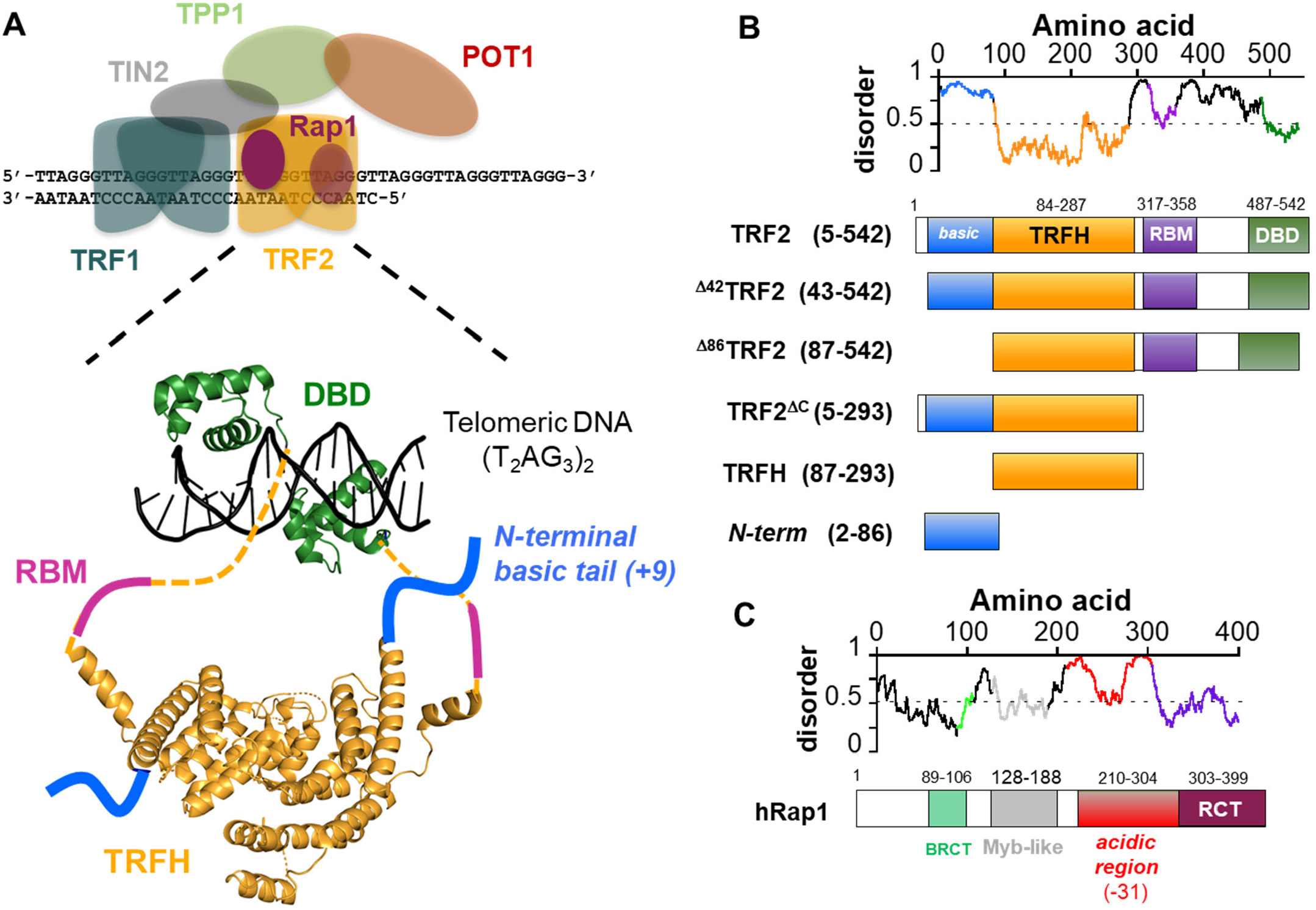
TRF2 and Rap1. **A**. Schematic representation of the 6 protein shelterin complex (TRF1, TRF2, Rap1, TIN2, TPP1, POT1). Enlarged is a representation of the architecture of TRF2 dimer bound to telomeric DNA: DBD, DNA-binding domain; TRFH, homodimerization domain; RBM, Rap1 binding motif. The dashed lines represent unstructured linkers connecting the TRFH to the DBD. Structures are based on 1h6p and 1w0u. **B**. Disorder/order predictions of TRF2 (IUPred) and schematics of the TRF2 constructs studied in this work. **C**. Schematic of Rap1: BRCT, BRCA1 C-terminal domain; Myb domain; acidic region with a −31 net charge spanning 100 aa; RCT, Rap1 C-terminus domain that interacts with the RBM of TRF2.

While telomeres are usually thought of as the nucleoprotein complexes formed at the ends of individual chromosomes, they are not always observed as distinct and separate nuclear entities. Though usually restricted in their motion, telomeres can transiently cluster together (40, 41) and dysfunctional telomeres assemble into telomere dysfunction-induced foci (TIFs) (42). Also, 10-15 % of cancer cells maintain their telomere length via Alternative Lengthening of Telomeres (ALT) (43, 44), a telomerase-independent pathway. In these cells, multiple telomeres co-localize with a subset of promyelocytic leukemia (PML) nuclear bodies to form ALT-associated promyelocytic leukemia bodies (APBs), within which telomeres are elongated (45-48). Intriguingly, PML bodies are a type of biomolecular condensates (14, 49), and a recent study has proposed that within APBs telomeres behave as phase-separated components (50). Whether the apparent phase-separated nature of APBs originates from an intrinsic ability of PML bodies to form biomolecular condensates or from one or more of the shelterin components to induce phase separation with telomeric DNA remains to be determined.

Of the two telomeric proteins that directly interact with DNA, TRF2 has long been implicated in DNA condensation *in vitro*, as monitored by gel during electrophoresis (51) or DREEM imaging (52). The molecular nature of these TRF2-DNA condensed states remains elusive. Single-molecule experiments (53) showed that pre-formed shelterin complexes do not lead to compaction of individual DNA chains or interaction between different DNA chains, suggesting that any ability of TRF2 to collapse/condense DNA may be modulated by interaction with other shelterin components. Also, individual telomeres typically appear as distinct globules of finite size, albeit T-loops where the 3’ ssDNA end is tucked into the dsDNA telomeric region can be visualized *in vivo* with high resolution imaging (54). These globules seem to be more compact than expected for equivalent lengths of bare dsDNA (55, 56), suggesting that interaction of one or more shelterin component with DNA may favor condensation of telomeres. However, studies testing the idea that decompaction of telomeres may be a means to regulate telomere accessibility *in vivo* reached divergent conclusions (57-59).

These observations raise key and unanswered questions – do telomeric proteins induce compaction of telomeric DNA? – if so, what are the physical and molecular bases for regulation of telomere compaction? – does phase separation play a role in establishing and regulating telomeres? To begin addressing these questions, we employed a collection of biophysical methods to directly test whether specific domains of TRF2 (Fig. 1B) induce the compaction of individual DNA chains and to investigate if and how binding of TRF2 to DNA leads to liquid-liquid mixing of the solutions. Our results reveal a symmetry between single-chain compaction and multi-chain condensation (*i*.*e*., phase separation), where perturbations that influence single-chain compaction are mirrored by perturbation that alters phase separation (such as salt concentration, binding of hRap1 to TRF2). Taken together, our results support the hypothesis that these emergent phenomena can be interpreted as two distinct outcomes controlled by the same set of intra- and inter-molecular interactions as well as by co-solutes.

## Results

### TRF2 promotes single-chain dsDNA condensation

We first sought to establish if TRF2 can drive single-chain DNA compaction. TRF2 has a modular architecture with an N-terminal basic disordered region, a central TRFH homo-dimerization domain, a disordered linker, and a C-terminal DNA-binding domain (DBD) that specifically recognizes T_2_AG_3_ telomeric repeats (Fig. 1A). When purified homo-dimeric TRF2 (Fig. S2A-C) is combined with telomeric repeat DNA, electrophoretic mobility into the gel is impeded in a DNA-dependent manner (Fig. S2D) (51), consistent with protein-mediated DNA condensation or aggregation. To directly test whether TRF2 compacts individual DNA chains, we employed two different single-molecule approaches.

First, we used two independently steerable optical traps and a laminar flow chamber to trap a single *λ*−DNA molecule in a dumb-bell configuration between two polystyrene beads (Fig. 2A, and top panel in 2B). By turning off the downstream trap relative to the flow, we varied the drag force on the free bead (*i*.*e*., the bead on the right in Fig. 2B) by varying the flow and measured DNA extension in the presence and absence of TRF2. In buffer alone, the DNA extension responds reversibly to changes in drag force (Fig. 2C, black trace). In contrast, upon movement of the DNA into a channel containing TRF2, a dramatic reduction in DNA extension was observed when the drag force was lowered. This DNA compaction was rapid and complete, as the free bead was eventually brought all the way into the active trap, thus ending the experiment (Fig. 2B and 2C). As expected for a protein-dependent phenomenon, the rate of DNA compaction increases with increasing TRF2 concentration (Fig. 2C).

**Figure 2.**
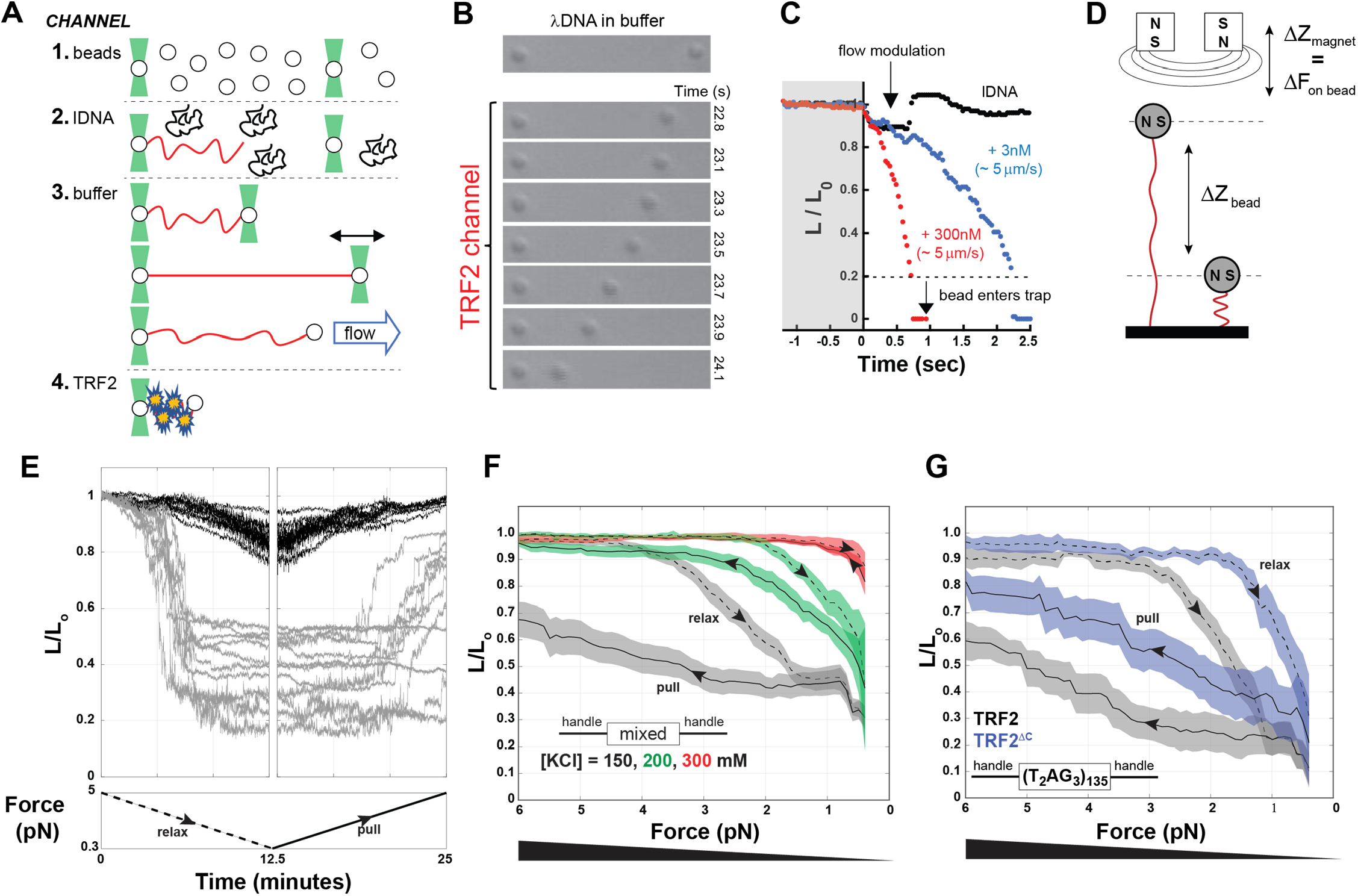
TRF2 condensation of single nucleic acids chains. **A**. Schematic of optical tweezers experiments. 1 to 4 are schematics of the laminar flows used to assemble *λ*DNA in a dumbbell configuration between two beads, before incubating it with protein. **B and C**. Compaction of DNA (expressed as the change in relative length L/L_0_) in presence of 300 nM (monomer) TRF2 as visually monitored by the decreasing distance between the beads over time (B) or quantified by tracking the position of the right bead (C), in the presence of TRF2 at 3 nM (blue) or at 300 nM (red) compared to the absence of TRF2 (black). **D**. Schematic of the magnetic tweezer experiment setup. **E**. Example of magnetic tweezer trajectories after relaxing (gray, first panel) and pulling (gray, second panel) the nucleic acid in the presence of TRF2. The DNA alone is show as reference in black. **F**. Quantification of DNA compaction as function of force for dsDNA of mixed sequence in the presence of 300 nM TRF2 (dashed and solid lines are the average and color bands the standard deviation of multiple traces in either the relax or pull direction, respectively). Increasing KCl concentration decreases the hysteresis between relax and pull experiments as well as the degree of force required to expand the conformations of the DNA. **G**. DNA compaction measured for the specific telomeric DNA sequence in presence of 300 nM TRF2 or its variant TRF2^ΔC^, which lacks the DBD and the linker region.

To further examine the force-dependence of TRF2-dependent DNA compaction, we turned to a magnetic tweezers assay which offered the benefits of higher throughput and the ability to precisely and stably establish constant forces between 0.5 - 5 pN. In these experiments, dsDNA molecules of ∼2 kbp effective length were attached at one end to a glass coverslip and at the other to a paramagnetic bead (Fig. 2D). DNA tethers were initially held at > 5 pN of force, the force was ramped down to < 0.5 pN, and then back to high force, with ramping at a rate of ∼ 0.6 pN / min (Fig. 2E). The force dependence of the end-to-end distance of bare DNA behaves as expected (60, 61) (Fig. 2E, black traces). Consistent with the optical tweezer experiments, the addition of TRF2 leads to DNA compaction (Fig. 2E, gray traces, first panel). We note that, at difference with bare DNA, in the presence of TRF2 the individual DNA tethers show a wide range of behaviors, including cases in which they remained irreversibly compacted even at the highest forces we applied (∼ 5 pN) (Fig. 2E, gray traces, second panel). This is suggestive of a highly cooperative mode of DNA condensation. Indeed, the presence of TRF2 leads to a significant hysteresis between relaxation (*i*.*e*., lowering the force) and pulling (*i*.*e*., increasing the force) (Fig. 2F), consistent with a large degree of cooperativity in the compacted state. TRF2-dependent DNA compaction and hysteresis were observed even at salt concentrations typically considered to be in the physiological range (Fig. 2F) and also with DNA tethers containing 0.8 kbp of human telomeric repeats ((T_2_AG_3_)_135_) (Fig. 2G in black).

Even though the biological function of TRF2 lies in its ability to specifically interact with telomeric DNA, our results clearly revealed that DNA compaction can occur independently of telomeric repeats. Since the DNA binding domain is known to interact specifically with telomeric repeats and these repeats appeared dispensable for compaction, we asked if the DNA binding domain (DBD) of TRF2 was also dispensable for DNA compaction. To our surprise, a TRF2 construct comprised of only the basic N-terminus and the dimerization domain (TRF2^ΔC^, Fig. 1B) can also condense single DNA chains (Fig. 2G in blue), albeit at lower forces and with less hysteresis, indicative of a more weakly-interacting system. These results suggest that the intrinsically disordered N-terminal domain directly contributes to DNA compaction, albeit perhaps in a non-specific manner.

To complement our single-molecule experiments we performed simulations using a hyper coarse-grained model in which a force constant is used to extend a single “DNA” chain while “TRF2” molecules interact with the chain in a multivalent manner (Fig. 3A). A first set of simulations was performed using two different TRF2:DNA interaction strengths (Fig. 3B, top). Importantly, TRF2:TRF2 interactions were completely absent, mimicking a scenario in where TRF2 does not engage in homotypic interactions. Under this regime, we observed a continuous transition in DNA extension as a function of force (Fig. 3B top, Fig. 3C). Such a result is in poor agreement with our force experiments (Fig. 2E), which do not show a continuous dependence on extension with force. Instead, force experiments appear to reveal two regimes; a largely extended DNA molecule (observed under high force) and a largely compact DNA molecule (observed under low/no force).

**Figure 3.**
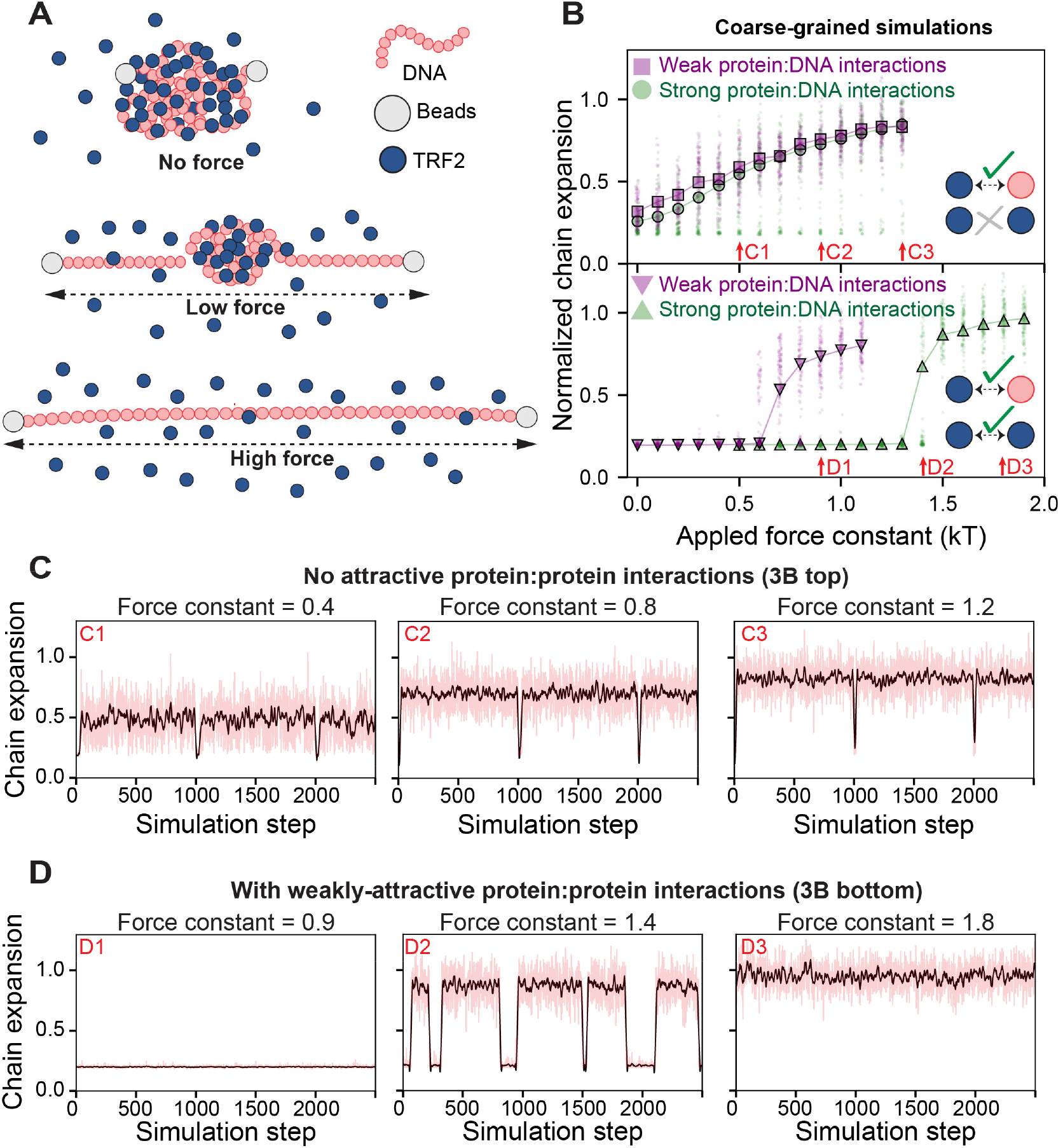
Simulation of TRF2-dependent DNA condensation. **A**. Schematic of setup for coarse-grained Monte Carlo simulations. A single DNA molecule of 80 beads was simulated with 4000 single-bead TRF2 molecules. The DNA end-to-end distance was constrained via a harmonic potential applied to the ends of the DNA (beads) with a variable force constant, mimicking force experiments. Under no force, TF2 molecules lead to DNA condensation, forming a spherical globule. Under low force, parts of the DNA are extended with a single spherical globule of DNA and TRF2 in the center. Under high force, the DNA molecule is fully extended with no TRF2 bound. **B**. Simulation results depend on the presence or absence of weak homotypic TRF2 interactions. If TRF2:TRF2 interactions are set to zero (*i*.*e*., implying TRF2 is infinitely soluble) we observed a continuous increase in chain dimensions as a function of force constant, both under weak (purple squares) and strong (green circles) protein-DNA interactions (top). If weak TRF2:TRF2 interactions are included (leading to a solubility limit of ∼0.5 mM for TRF2), we observe apparent two-state behavior (bottom). Here the DNA is either compact or expanded, as characterized by the sharp transition as a function of force and the co-existing populations giving rise to a bimodal distribution. The force constant at which expansion occurs depends on the strength of the protein:DNA interaction. Arrows reference simulation traces in panels 3C and 3D. **C**. Individual simulation traces for simulations performed absent TRF2:TRF2 interactions. Red lines are true data, black line is a smoothed trace. Occasional troughs are transitions into short-lived metastable compact globular states. **D**. Individual simulation traces for simulations performed with weak TRF2:TRF2 interactions, revealing two-state behavior. During a single simulation, the system oscillates between two fixed points (compact and expanded). This oscillation is analogous to an infinitely cooperative transition, as seen when sitting at a phase boundary. Red lines are true data, black line is a smoothed trace.

Why does our model not recapitulate the gross behavior of the force experiments? One possibility is the limiting assumption that TRF2 cannot engage in any kind of attractive interactions with itself. While TRF2 is monodisperse in isolation (Fig. S1B), it contains multiple disordered regions which may engage in both specific and non-specific protein-protein interactions (62). More generally, proteins are typically not infinitely soluble under aqueous solutions, implying the presence of weak, non-specific intermolecular interactions (63). With this in mind, we considered an alternative model in which weakly attractive TRF2:TRF2 interactions were included. These interactions are extremely weak; in the absence of DNA, the protein remains soluble up to around ∼0.5 mM (∼30 mg/ml, 3x more soluble than lysozyme). Despite this, the inclusion of weak TRF2:TRF2 interactions fundamentally shifts the dependence of DNA compaction on force (Fig 3B, bottom). In simulations performed with weak TRF2:TRF2 interactions, we observe a highly cooperative transition where the chain is *either* fully compact *or* fully extended, in line with the observations in force experiments (Fig. 2E, Fig. 3B and 3D). Importantly, even within a single simulation apparent two-state behaviour emerges, with the system jumping between a compact globule and an expanded chain (Fig. 3D).

Taken together, both experimental and simulated data strongly suggest that while the DBD of TRF2 contributes to an increase of interaction strength due to the specific binding to the T2AG3 repeats, weaker protein-DNA interactions in the context of the multivalent nature of TRF2 may be key drivers of the cooperative compaction of single DNA chains.

### Interaction of TRF2 with multiple dsDNA molecules leads to phase separation

Polymer theory predicts that the same molecular interactions that lead to collapse of a single DNA chain can also lead to condensation when multiple chains are present in solution at high enough concentration (18). Indeed, TRF2 forms complexes with DNA that during electrophoresis barely migrate into the gel (Fig. S1D) (51), consistent with formation of either large supramolecular assemblies, amorphous aggregates or, possibly, phase separation. With these possibilities in mind, we found that while TRF2 is a well-behaved dimer in solution (Fig. S1), at a sufficiently high concentration mixing of TRF2 with non-specific dsDNA (supercoiled or linear fragments) results in an opalescent solution that contains micron-sized droplets, as revealed by Differential Interference Contrast (DIC) microscopy (Fig. 4A and S3A). In addition, these droplets fuse over time (Fig. S3C), consistent with a system that has undergone phase separation to form a dynamic and liquid-like dense-phase.

**Figure 4.**
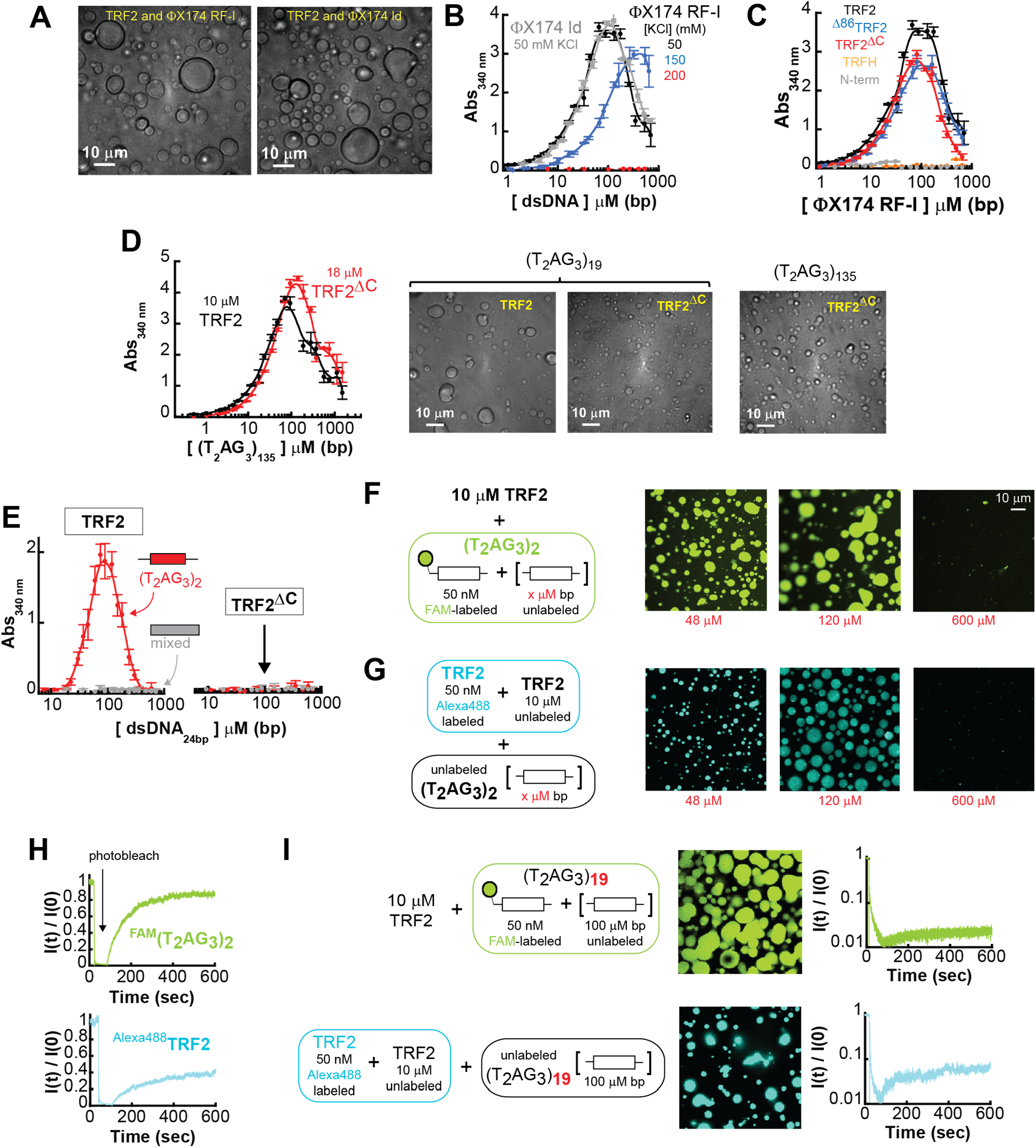
TRF2 phase separation with specific and non-specific dsDNA. **A**. DIC microscopy images of phase-separated droplets in presence of ϕX174 RF-I at the indicated KCl concentration, and ϕX174 HaeIII digest (gray). **B**. Turbidity measurement of solutions containing a fixed concentration of TRF2 (10 μM monomer) and variable concentrations of ϕX174 RF-I (black) or ϕX174 ld (gray). Increasing salt concentration from 50 mM KCl to 150 mM KCl (blue) shifts phase boundaries to higher concentrations. A further increase to 200 mM KCl (red) suppresses phase separation at this concentration of TRF2. Error bars represent the standard deviation from three repeats, and the solid lines are interpolations of the data to facilitate visualization. **C**. Turbidity experiments of solution containing non-specific ϕX174 RF-I in presence of truncation variants of the protein at fixed concentration of 10 μM monomer. Turbidity is suppressed for the TRFH construct and strongly reduced for the N-terminal tail fragment. **D**. Turbidity measurements of TRF2 and TRF2^ΔC^ variant reveals phase separation in presence of specific ds-DNA sequences. **E**. TRF2 (10 μM mon) undergoes phase separation only with a 24 bp DNA that contains a (T_2_AG_3_)_2_ repeat in its center, and not with a DNA of mixed sequence composition of the same length. Under identical conditions truncation of the C-terminus (TRF2^ΔC^) suppresses phase separation also for specific DNA. **F-G**. Re-entrant behavior of the phase separation with increasing concentration of specific dsDNA. **H-I**. FRAP of labeled TRF2 and specific DNA mixtures shows order of magnitude decrease in mobility with increasing length (from 2 to 19) of the number of specific telomeric binding sites on the dsDNA substrate.

The formation of scattering objects (*i*.*e*., droplets) was further assessed by absorbance scattering at 340 nm, where neither protein nor nucleic acid absorb. The solution absorbance displayed classical re-entrant behavior as a function of DNA concentration (64) (Fig. 4B). Absorbance initially increases, reflecting the formation of droplets; however, as the DNA concentration is increased further, absorbance then decreases, in line with the ‘re-entrance’ into the one-phase regime. This re-entrant behavior is expected for a two-component system governed by a closed-loop phase-diagram (64). Whereas the re-entrant phase may be interpreted in the context of charge inversion of oppositely charged polyelectrolyte (65), the phenomenon can occur also when the interaction is realized between uncharged domains (66). More in general, in a mixture of two different components, the re-entrant phase reflects the saturation of the binding sites of one component by the second component, such that binding across two different molecules of the same component is disfavored.

Phase separation occurs over the same concentration regime with either supercoiled DNA or its linear fragments, and at a physiological range of salt concentrations (Fig. 4B). Importantly, the salt concentrations over which phase separation is observed mirror those under which TRF2-dependent DNA compaction occurs. Taken together, our results demonstrate that TRF2 can induce both DNA compaction and phase separation.

Having established that TRF2 can drive phase separation with DNA, we designed a series of TRF2 constructs to assess the contribution of specific protein regions to phase separation (Fig. 1B and S1A-C). Neither deletion of the basic domain (^Δ86^TRF2) nor removal of the DBD and the unstructured linker (TRF2^ΔC^) suppressed DNA-dependent phase separation (Fig. 4C). These observations indicate that TRF2-dependent phase separation is neither dictated exclusively by the presence of the disordered N-terminal basic region nor by the linker and DBD. Moreover, neither the TRFH domain nor the N-terminal region alone promote phase separation at the same protein concentrations as observed for full length protein (Fig 4C) but can be driven to phase separate with DNA at higher concentrations (Fig. S3D). These data strongly suggest that multivalent interactions with the nucleic acids are required for efficient phase separation and that multi-valency is realized *via* both unstructured and structured domains of TRF2 that act in a synergistic fashion.

### Specificity and multi-valency of telomeric repeats

Given the specificity and high affinity with which the DBD of TRF2 binds telomeric repeat sequences (67), we would anticipate that at cellular concentrations, telomeric repeat DNA will be preferentially bound by TRF2. With this in mind and considering TRF2 ability to undergo phase separation with non-specific dsDNA, we anticipated that TRF2 would also phase separate when mixed with telomeric repeat sequence DNA. However, in contrast to non-specific dsDNA, we expected that TRF2 phase separation with telomeric repeat DNA would be strongly dependent on the presence of the DNA binding domain. As expected, solution demixing was observed in the presence of telomeric repeat-containing DNA and TRF2 (Fig. 4D). Yet, while slightly higher protein concentrations are required to achieve equivalent turbidity, the TRF2^ΔC^ construct, which lacks the DBD, is still able to drive DNA-dependent phase separation with dsDNA that contains multiple telomeric repeats (Fig. 4D). Taken at face value, these data could be interpreted to suggest that DNA-dependent phase separation is driven by non-specific interactions of TRF2 with DNA, with the specific recognition of telomeric repeats by the DBD playing a limited role in the process. Alternatively, the multivalent interactions mediated by different TRF2 domains and the DNA may mask the impact of DNA specificity encoded by the DBD in the context of telomeric repeat sequences.

To delineate between these two interpretations, we sought to test for the presence of DNA-specific interactions with shorter DNA repeat sequences. For this, we turned to a short dsDNA fragment of 24 bps that contains a single (T_2_AG_3_)_2_ repeat in its center and a length-matched control that lacks the (T_2_AG_3_)_2_ repeat (mix sequence dsDNA). Strikingly, at the protein concentration used in the assay, phase separation occurs only in the presence of telomeric repeats (Fig. 4E). Furthermore, independent of the presence of repeats, a significant suppression of phase separation was observed with the TRF2^ΔC^ construct that lacks the DBD (Fig. 4E), in stark contrast to what was observed with longer DNA substrates. Thus, once the valence of the nucleic acid is reduced by decreasing its length, the importance of DBD-encoded specificity for telomeric repeats is clearly unmasked.

The importance of DBD-mediated specific interaction is further supported by fluorescence confocal imaging experiments. Droplets of TRF2 and a fluorescein-labeled 24 bp dsDNA containing the (T_2_AG_3_)_2_ repeat, formed in the presence of different concentrations of unlabeled DNA (Fig. 4F), recapitulate the absorbance scattering experiments. As the concentration of DNA increases larger and more numerous droplets form, followed by their disappearance at high DNA concentration as expected for a re-entrant phase. Similar results are obtained when Alexa-488 labeled TRF2 is used in place of labeled DNA (Fig. 4G). In addition, Fluorescence Recovery After Photobleaching (FRAP) of the fluorescent droplets indicates that a large fraction of the DNA within the droplet remains mobile (Fig. 4H), consistent with a liquid-like character of the dense phase. Furthermore, TRF2 inside the droplets is mobile as well, albeit to a lesser extent than DNA. Droplet liquidity is maintained for at least 24 hours (Fig. S4A), suggesting that if further molecular re-arrangements lead to solidification (*e*.*g*., fibril formation), this occurs at significantly longer timescales (69, 70). In stark contrast, TRF2 form significantly fewer droplets with the 24 bp dsDNA of mixed sequence composition (Fig. S4B) and TRF2^ΔC^ shows a further reduction in droplets independently of the nature of the nucleic acid sequence (with or without telomeric repeat, Fig. S4C and S4D), consistent with the results of the turbidity experiments.

Therefore, our data support a model in which the multivalency of both the protein and the DNA dictate the phase separation propensity of the system. Once the valence of the DNA is reduced by shortening its length, non-specific TRF2-DNA interactions are no longer sufficient to drive phase separation and the specific recognition of a telomeric repeat by the DBD becomes essential. On the other hand, this model also predicts that as the valence of the DNA increases TRF2 should phase separate in a seemingly non-specific manner even in the absence of its DBD. Indeed, consistent with the absorbance scattering and DIC experiments in Figure 4D, droplets are also observed with a dsDNA that contains 19 T_2_AG_3_ repeats (Fig. 4I), independent of the presence of the DNA binding domain (Fig. S4E). Interestingly, the droplets formed by TRF2 and the (T_2_AG_3_)_19_ telomeric substrate show limited recovery after photo-bleaching, independent of whether the DNA or TRF2 are being monitored (Fig. 4I). This behavior is different to that observed from the 24 bp dsDNA and two T_2_AG_3_ repeats and it suggests that the increase in valence with multiple telomeric repeats leads to a dramatic change in the physical properties of the phase separated state. While these droplets show slow internal dynamics, they are still able to fuse over long timescales (Fig. S5).

### Phase separation with single-stranded nucleic acids

Transcription of telomeric repeats leads to telomeric repeat-containing RNA (TERRA), a long non-coding RNA proposed to play a role in telomeric protection (71, 72). The interaction between TERRA and TRF2 has been proposed to facilitate heterochromatin formation, while TERRA depletion leads to an increase in TIFs (73). Intriguingly, a splice-variant of TRF2 lacking the linker and DBD (similar to TRF2^ΔC^) co-localizes with neuronal granules, ribonuclear biomolecular condensates that facilitate the transport of mRNAs in axons (74, 75). Based on our work here, one interpretation of this observation is that while the shorter TRF2 splice variant loses its ability to interact with telomeres it retains non-specific binding affinity for RNA, possibly via interactions within its basic N-terminus domain. With this in mind, we wondered if the DNA-dependent phase separation of TRF2 was truly dependent on dsDNA, or if phase separation could also be driven and modulated by other types of single-stranded nucleic acids (ssNAs), as has been shown for other non-specific positively charged polypeptides (76).

In line with this expectation, absorbance scattering experiments and DIC imaging indicate that TRF2 phase separation occurs with both ssDNA (Fig. 5A) and ssRNA (Fig. 5C) over a similar range of salt concentrations as it did with dsDNA (Fig. 4B). As with dsDNA, this phase separation requires only the N-terminal basic domain or the DBD (Fig. 5B and 5D). However, in contrast to the lack of phase separation observed with the short 24 bp DNA of mixed sequence composition, at the same protein concentration non-specific interaction of TRF2 with an oligo-dT ssDNA as short as 20 nt is sufficient to induce phase separation (Fig. S6A). An interpretation of this result is that, because of chain flexibility, non-specific ssDNA interactions occur via a larger number of contacts than non-specific dsDNA interactions, increasing the protein-nucleic acid affinity and lowering the critical concentration for phase separation. In summary, our results demonstrate that TRF2 can phase separate with a variety of nucleic acids, where the driving force for phase separation depends on the strength and valence of interactions encoded by both proteins and nucleic acids.

**Figure 5.**
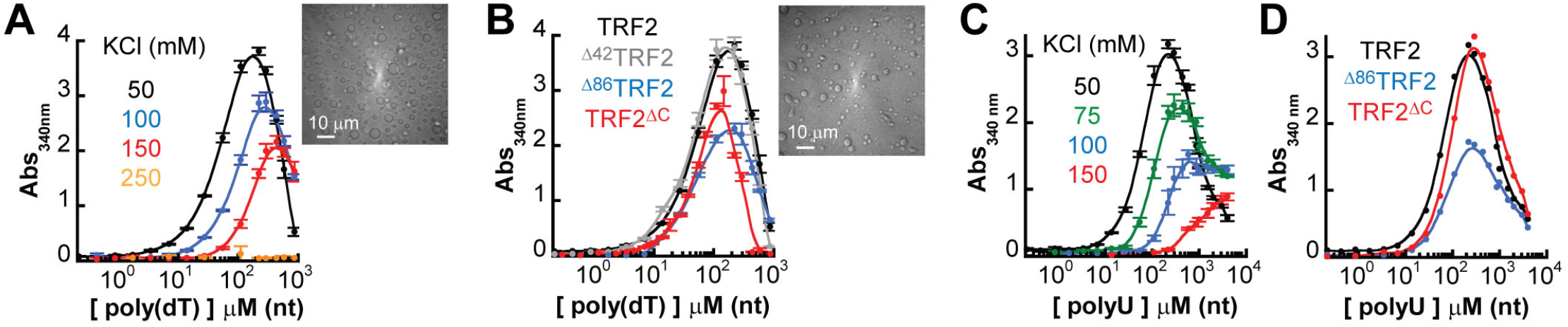
TRF2 undergoes phase separation with ssDNA and ssRNA. **A**. Turbidity experiments with mixtures of full length TRF2 (10 μM mon) and poly(dT) at increasing concentration of KCl concentration reveals a shift of the saturation concentration toward higher values and suppression of demixing at 250 mM KCl. **B**. Phase separation occurs also in truncated variants of the protein that lack the first 42 residues (Δ42), the first 86 residues (Δ86) or the last 294-542 amino acids (ΔC). **C**. Turbidity experiments with mixtures of full length TRF2 (10 μM mon) and poly-rU reveals a shift of the saturation concentration toward higher values with increasing salt as for ssDNA. **D**. Truncation variants of the protein maintain the phase separation propensity of the protein with poly-rU.

### hRap1 modulates single DNA chain condensation and the phase-separation propensity of TRF2

We have described the high propensity of TRF2-dependent DNA condensation and phase separation. However, given that TRF2 is only one component of the shelterin complex, we next asked how these processes might be modulated by the presence of a binding partner. More specifically, within the core-complex (TRF2-hRap1-Tin2-TRF1), hRap1 is unique in that it interacts directly and exclusively with TRF2 (29, 33, 34), making hRap1 a clear partner to test. *In vitro*, TRF2 and hRap1 form a tight complex (33) and, at the protein concentrations used in our experiments (> 300 nM), the TRF2-hRap1 complex can be considered as a single component. Binding of hRap1 to TRF2 lowers the DNA binding affinity of TRF2 (33). Moreover, the TRF2 interacting domain of hRap1 (RCT in Fig. 1D) is preceded by a highly acidic region that may interfere with the function of the basic domain of TRF2. Thus, it is reasonable to postulate that interaction of hRap1 with TRF2 may modulate both the TRF2-dependent compaction of a single DNA chain and the DNA-dependent phase-separation propensity of TRF2. Indeed, magnetic tweezers experiments indicate that, compared to TRF2, the TRF2-hRap1 complex induces a smaller extent of compaction of DNA tethers containing the telomeric repeats and a lower degree of hysteresis (Fig. 6A). These observations are consistent with hRap1 reducing the affinity of TRF2 for DNA (33), effectively altering the valence of TRF2, and modulating the properties of the collapsed state.

**Figure 6.**
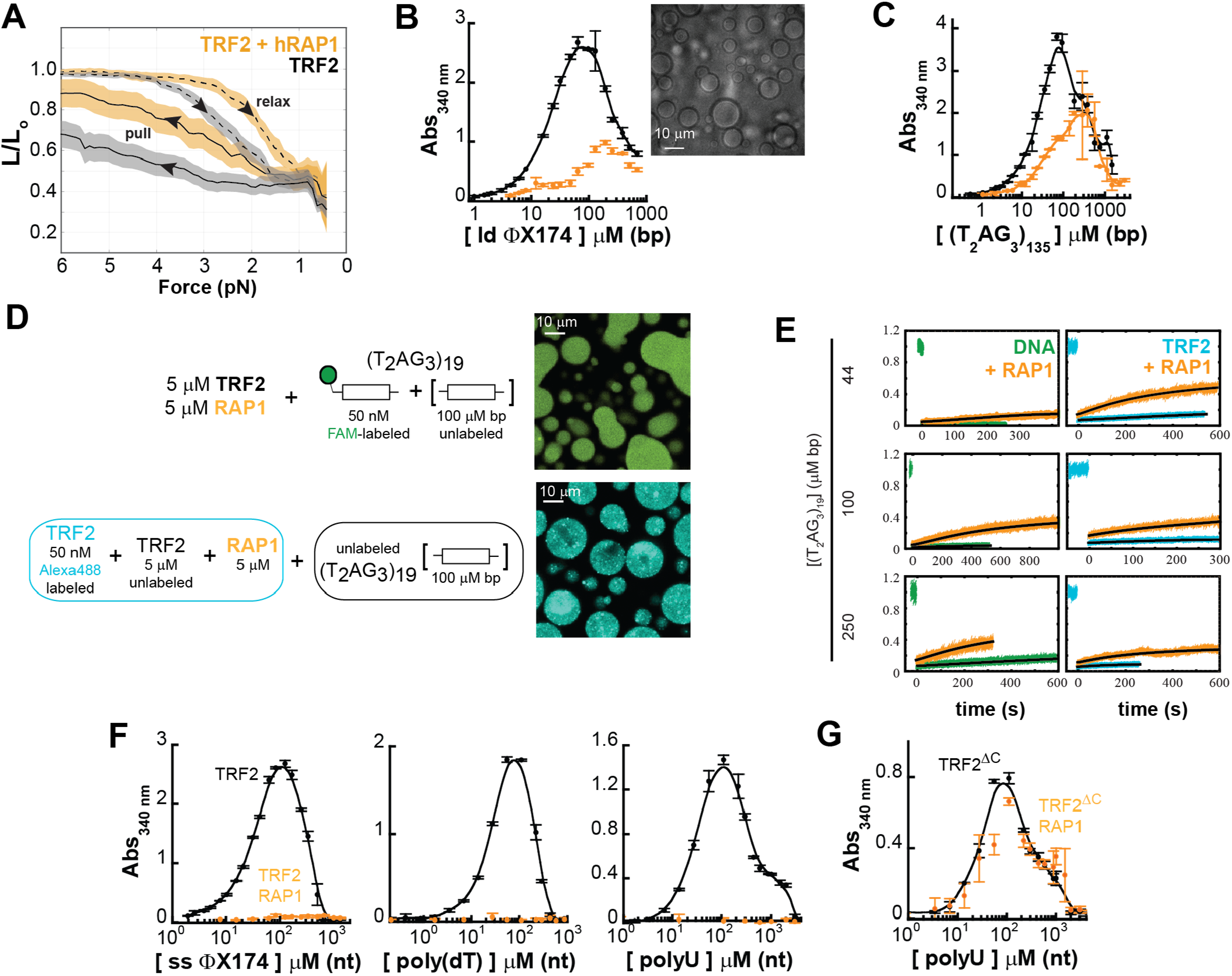
hRap1 modulates the phase-separation propensity of TRF2 with ds and ss nucleic acids. **A**. Force-extension experiments reveal a smaller TRF2-dependent DNA compaction in presence of stoichiometric concentration of hRap1 (dashed and solid lines are the average and color bands the standard deviation of multiple traces in either the relax or pull direction, respectively). **B-C**. Turbidity measurements of TRF2 and a TRF2-hRap1 stoichiometric complex in presence of non-specific (ϕX174 HaeIII digest) and specific (T_2_AG_3_)_135_ double-stranded DNA. Addition of hRap1 does not ablate phase separation but modulates the phase-boundaries. **D-E** Phase separation of TRF2 (5 μM mon) and hRap1 (5 μM) with (T_2_AG_3_)_19._ Alexa488-labeled TRF2 and FAM-labeled DNA demonstrate partitioning of the protein and DNA in the dense phase. FRAP measurements reveal that addition of hRap1 increases under all conditions the recovery of both DNA and TRF2. **F**. Turbidity measurements of TRF2 in presence of ss ϕX174, poly(dT), and poly(U) reveal suppression of phase separation upon addition of stoichiometric concentrations of hRap1. **G**. Turbidity measurements of the truncation variant TRF2^ΔC^, which lacks the binding site of hRap1. No change in turbidity is observed upon addition of hRap1.

Although TRF2-hRap1 interaction with non-specific double-stranded DNA still leads to formation of large micron-size droplets (Fig. 6B and S3A), the change in absorbance scattering in the presence of hRap1 suggest that the phase boundaries have shifted. While binding of hRap1 to TRF2 does not suppress phase separation with dsDNA that contains telomeric repeats, there is a diminution in the driving force for assembly, mirroring changes observed in the force experiments (Fig. 6C, 6D, and S3B). Interestingly, the presence of hRap1 leads to an increase in the recovery after photobleaching of the labeled DNA in the droplets (Fig. 6E), consistent with its ability to reduce the strength of TRF2-DNA binding.

Finally, at the concentrations used in our experiments, interaction of hRap1 with TRF2 abolishes (or shifts to significantly higher concentrations) phase separation with single-stranded nucleic acid, either DNA or RNA (Fig. 6F). These results suggests that hRap1 may function as a specificity switch for different types of nucleic acids, reducing non-specific DNA binding more severely than specific (telomeric repeat) DNA binding. It is important to note that the effect of hRap1 is mediated by its direct interaction with TRF2, as hRap1 has no effect on the phase separation of the truncated variant TRF2^ΔC^ that lacks the interaction region with hRap1 (Fig. 6G).

If TRF2 undergoes *bona fide* phase separation with dsDNA, then it should be possible to construct a complete two-dimensional phase diagrams with a closed-loop binodal, as predicted by analytical theories of polymer mixtures (77-80). Moreover, if hRap1 weakens DNA compaction and phase separation by modulating protein-DNA interaction then that phase diagram should be systematically tuned by the presence of hRap1, shifting towards higher saturation concentrations (81). At the same time, binding of hRap1 to TRF2 will introduce further excluded volume in the dense phase and therefore cause a shift toward lower concentrations of TRF2 in the dense phase.

In support of this hypothesis, we were able to infer the close-loop nature of the two-dimensional phase diagram for TRF2 with dsDNA by measuring the concentrations of TRF2 and nucleic acids in the light and dense phase (Fig. 7A). Measurements are carried out using 50 nM of labeled protein or DNA mixed with excess unlabeled protein and nucleic acid (at a ratio of at least 1:200, see Supplementary Information). The use of single-photon detectors enables the determination of the concentration in the light phase with sensitivity down to the pM concentration and limits artifacts in the determination of protein and nucleic acid concentrations in the dense phase, which could arise at high concentration of fluorescent species due to quenching of the fluorophore or saturation of the detectors. The linearity range of the approach has been previously validated for a single component phase-separation system, by comparing the concentration obtained by fluorescence intensity with those measured with fluorescence correlation spectroscopy as well as independent turbidity measurements (28). Our measurements provide an estimate for the directionality of the tie-lines, which support the notion that protein and nucleic acid favorably co-phase separate, as indicated by the positive slope in the protein-nucleic acid space. When comparing the concentration at which phase separation occurs, variation of the length of the DNA segment systematically alters the phase boundaries, as predicted by theory and in strong agreement with a three-component (solvent, DNA, protein) phase diagram (Fig. 7B). Also, upon addition of hRap1, the phase boundary relative to the saturation concentrations recesses toward higher values, reporting on a systematic diminution in protein-DNA interaction driven by hRap1, as anticipated based on the force experiments (Fig. 6B).

**Figure 7.**
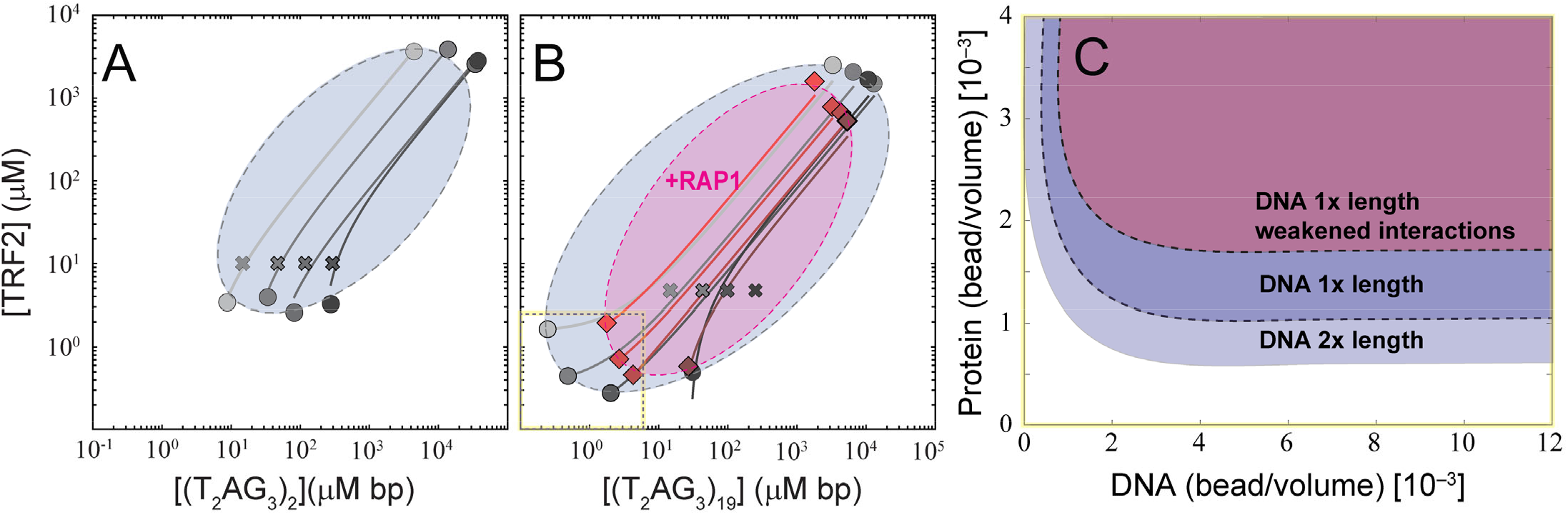
TRF2-DNA phase diagram and the implications for multi- and single-DNA condensation. **A-B**. Reconstruction of the phase diagram for mixtures of TRF2 and (T_2_AG_3_)_2_ (A) and TRF2 and (T_2_AG_3_)_19_ (B) in gray as obtained by measuring the intensity of labeled protein and DNA in the dilute and dense phase (full circles) after mixing of the indicated concentrations (crosses). Solid lines connecting the dilute and dense phase concentrations represent expected tie lines across the solution, whereas the dashed lines are guide-to-the-eyes phase boundaries based on the concentration limits determined *via* fluorescence intensities and turbidity measurements. Addition of Rap1 (magenta area) restricts the phase boundaries. **C**. Lattice-based multi-component simulations reproduce the observed change in boundaries when increasing length of the DNA or decreasing multivalence of the TRF2 through a mean-field diminution in attractive interactions, mimicking the interpreted effect of hRap1. Only the lower-left portion of the phase diagram in panel B is simulated in panel C (dashed line with yellow highlight).

The phase behavior observed in experiment is recapitulated in simple coarse-grained simulations. Longer DNA molecules lead to an increased propensity to undergo phase separation as reported on by the reduction in saturation concentration (Fig 7C). Moreover, when we weakened intermolecular interactions in the same manner as in Fig. 2H and S7 (which leads to a decrease in compaction) we systematically shift the saturation concentration to higher values (Fig. 7C), analogous to the change observed upon addition of hRap1 (Fig. 6B).

Finally, the concentration of TRF2 in the dense phase is also affected by binding of hRap1, in agreement with our excluded volume argument (Fig. 7B). It is interesting to note that across the phase-diagram, addition of hRap1 alters the amplitude of FRAP (which is consistent with a weakening of interactions between TRF2 and nucleic acids), but it does not significantly affect the diffusion time of either DNA or TRF2 in the dense phase (Fig. S8). This observation is consistent with an increase in the fraction of the mobile species, where the mobility of the diffusive species is not largely dissimilar in the presence or absence of hRap1. Overall, these observations support the idea that the interaction of specific ligands (such as hRap1) can control (and even suppress) both phase separation (phase boundaries, selectivity for specific nucleic acids, transport properties) and nucleic acid compaction by modulating the set of interactions encoded by the binding partner (e.g., TRF2).

## Discussion

Here we showed that two of the shelterin components, TRF2 and hRap1, modulate both protein-protein and protein-DNA interactions and lead to either compaction of individual DNA chains or phase separation of proteins and DNA at higher DNA concentrations. Similar results can be recapitulated using coarse-grained simulations that capture the overall trends of DNA compaction and phase separation propensity in the presence of a ligand. Taken all together, our observations support the original interpretation proposed by Post and Zimm (18) and are consistent with the idea that protein-dependent collapse of a single nucleic acid chain and the phase separation of mixtures of protein and nucleic acids are two distinct outcomes driven by the same set of molecular interactions (68, 82, 83).

The multi-valence encoded in both proteins and nucleic acids is clearly key for creating a network of molecular interactions that can engage both intra-molecularly (single-chain condensation) and inter-molecularly (phase separation). More specifically, our experiments reveal that the modular architecture of TRF2 encodes for a certain redundancy, such that phase separation with nucleic acids is robust even in absence of either the N-terminal basic domain or the DBD itself. Droplets also form when DNA is assembled in poly-nucleosomes (Fig. S6B), suggesting that TRF2-dependent phase separation may occur with more complex protein-DNA assembly that are formed in cells. Indeed, several recent studies have proposed or demonstrated that telomeres function as dynamic liquid-like condensates in living cells (50, 84, 85). Given the repetitive nature of telomeric DNA, our observations suggest a model in which telomeric repeats provide a multivalent platform for the formation of dynamic, sheltrin-mediated condensates at telomeres, driving compaction of telomeric DNA. This model mirrors similar conclusions arrived at independently by the Yildiz and Brangwynne labs through a mixture of *in vitro* and *in cell* experiments (84).

Despite of the presence of a domain with high specificity, in our assays TRF2 is generally promiscuous in its interaction with nucleic acids, including specific and non-specific dsDNA as well as ssDNA and ssRNA, raising the question of how these types of interactions are regulated in cells. One possibility is that specificity is modulated by the presence or absence of additional binding partners. Along these lines, we find that hRap1 significantly alters TRF2’s propensity to both condense and phase-separate nucleic acids. Specifically, binding of hRap1 to TRF2 selectively abolishes phase separation with ssNAs, thereby imparting a dsDNA substrate specificity for the full-length protein. Deletion of the TRF2 region where hRap1 binds, which mimics the natural splicing form interacting with RNA, restores phase separation with single-stranded nucleic acids and suppresses specificity for dsDNA. In other words, the binding of hRap1 acts as a switch that controls the balance between promiscuous and specific interactions of TRF2 with nucleic acids. We propose that this mechanism may be important in the cell to discriminate TRF2 functions at telomeres versus the functions of its spliced variant in RNA transport in neurons.

Broadening this model, these observations suggest that perhaps different shelterin components may play a role in dictating the degree of collapse of single telomeres and favoring or disfavoring the phase separation with multiple nucleic acids. It is important to stress that though TRF2 undergoes phase separation with both non-specific and specific dsDNA, the specificity of telomeric DNA sequences and the number of specific telomeric sites have an impact on the transport properties of the phase-separated state, as monitored by FRAP. Therefore, specific sequences may have different mobilities and miscibilities, *de facto* contributing to the separation of different nucleic acids components.

To avoid ambiguity, we want to clarify that our results do not imply that individual telomeres are a *bona fide* phase-separated state. Rather, we propose that the mode of interaction between shelterin components and telomeric DNA determines whether single telomeres remain distinct entities or merge into larger bodies. Moreover, we propose that the interactions that drive *bona fide* phase separation in our *in vitro* reconstitution mirror the interactions that drive compaction of telomeric DNA, albeit perhaps with different affinities and specificities, as determined by the complement of binding partners and the local environment. This does not exclude the possibility that specific local regions of nucleic acid may favor a local partitioning of proteins, analogous to the pre-wetting demixing that occurs near to surfaces (86) and as recently shown for the interaction of FUS in proximity of stretched DNA (87). Furthermore, coalescence and assembly of the multiple telomeres or fragment of telomeres may explain the recent observation and proposal that telomeres in APBs have properties consistent with being a phase-separated state (50).

### Ideas and speculation

The phase-separation propensity of TRF2 and its modulation by interaction with hRap1 may be part of a more general mechanism for the regulation of nucleic acids in cells. Indeed, recent experimental evidence indicates that HP1a, a major component of heterochromatin, can also form liquid droplets and such a phenomenon is proposed to be the basis of heterochromatin organization (7, 8, 21). One interpretation of these results reflects a model in which HP1a drives heterochromatin territories through *bona fide* liquid-liquid phase separation (7, 8, 21). Another is that the intrinsically multivalent nature HP1a and many other DNA binding proteins (such as TRF2) drives DNA compaction through distributed multivalent interactions which will inevitably also drive phase separation with sufficiently short nucleic acid molecules *in vitro* (68, 82, 88). Altering the degree of DNA compaction by modulating the same interactions that would also drive phase separation could allow for the rheostatic control of access to genomic loci. These interactions may be modulated through subtle changes in the complement of available binding partners (*e*.*g*., with hRap1, as we show here), but also by chemical modifications to proteins or nucleic acids (*e*.*g*., histone acetylation or DNA methylation). Similarly, super-enhancers have been proposed to assemble through phase separation of transcription factors at specific locations in the genome (89-92). Another example is the recent finding that the nucleocapsid protein of the SARS-CoV-2, which plays a role in the viral RNA condensation and packaging, can also undergo phase separation and localize in biomolecular condensates within the cell (93-95). This observation has led to a proposed model in which viral genome compaction is driven by the same interactions that can also form biomolecular condensates (68). In this respect, the finding that telomeric components can lead to solution demixing supports the idea that the underlying physics of phase separation may be another example of a general mechanism through which the access and state of genetic material is controlled.

Our results highlight how phase behavior can be modulated by specific protein-ligand interactions, stressing the importance of investigating the contribution of linkage effects to phase separation (81). Indeed, our results suggest that at telomeres specific interactions within the shelterin complex may be a means to regulate and suppress the phase-separation propensity of single components, such as TRF2. Similar mechanisms may have evolved in nature to modulate the multivalence of a specific protein, regulating its phase-separation propensity, specificity, or selectivity. Finally, the overall picture emerging from the parallelism between single chain condensation and multiple chain phase separation, as originally proposed by Post and Zimm (18) (see SI Appendix and Fig. S9 for additional discussion on this aspect), suggests that phase separation can be used as a readout for the condensation of single chains and, as such, can provide a convenient macroscopic tool to investigate and test therapeutics (96, 97).

### Methods

A detailed description of the experimental and computational methods and approaches used in this work is provided in Supplementary Information.

## Data Availability Statement

All the data have been included in the figures or provided in the Supporting Information section.

## Acknowledgements

We would like to thank John Cooper for access to the microscope for DIC imaging. This work was supported by the National Institutes of Health 1R35GM139508 to R.G. and 1R01AG062837 to A.S.

