## Supplementary Information for "Shelterin components modulate nucleic acids condensation and phase separation in the context of telomeric DNA"

### Cloning and Site-Specific Mutagenesis.

All the primers sequenced used in this work are reported in Table S1. All the DNA and gene sequence used as templates were purchased from Addgene. Gene amplification was carried out using a two steps PCR protocol with Phusion DNA polymerase (NEB cat. M0530), while PfuTurbo polymerase (Agilent cat. 600250) was used for standard site-specific mutagenesis. Correct sequences for all the generated plasmids were confirmed by DNA sequencing.

*hTERF2 constructs* – All the gene amplification PCR reactions were carried out in GC buffer and in the presence of 1 M betaine. All the hTERF2 constructs were cloned at EcoRI/XhoI restriction site of the pGEX-6p-1 vector. Plasmid p16-1 hTRF2 (Addgene cat. 12299) was the template for the cloning of the full-length protein using primers hTERF2-F and hTERF2-R. In the resulting plasmid, the gene was not in frame with the GST and missing the $Ala^{477}$. Primers h-TERF2-frame-F, h-TERF2-frame-R, hTERF2-$Ala^{477}$-F and hTERF2-$Ala^{477}$-R were designed to take care of these issues. The final plasmid (pGEX-hTERF2) coded for TERF2 sequence spanning from residue 5 to residue 542 (isoform 1 id:Q15554-3) and it was used as templated for the generation of all the hTERF2 constructs. Plasmids pGEX-hTERF2$^{\Delta 42}$ and pGEX-hTERF2$^{\Delta 86}$ were generated by regular cloning using hTERF2Δ42-F and hTERF2Δ86-F respectively as forward primer, and hTERF2-R as reverse primer. The C-terminus deletion constructs and pGEX-hTERF2$^{\Delta C}$ was generated by site-specific mutagenesis using primers hTERF2-S294Stop-F and hTERF2-S294Stop-R. Primers hTERF2-S294Stop-F and hTERF2-S294Stop-R were also utilized for the generation of plasmid pGEX-hTERF2$^{\Delta 86\text{-}\Delta C,}$ yielding the only the TRFH domain of TRF2. Plasmid pGEX-hTERF2$^{(2\text{-}86)CC}$, encoding for sequence encompassing 2 and 86 residues and with the additional mutations S12C and A85C, was generated by PCR amplification of this sequence from a gBlocks-hTERF2-N-term (IDT, Corvallis) using primers hTERF2-A2-F and hTERF2-G86-R and cloning it at BamHI/XhoI site of pGEX-6p-1. Plasmid pGEX-hTERF2-$Cys^{160}$ expressing hTERF2 mono-cysteine mutant at position 160 was obtained by iterative site-specific mutagenesis using primers hTERF2-C148S-F, hTERF2-C148S-R, hTERF2-C207S-F and hTERF2-C207S-R.

*hRap1 construct* - Two steps were necessary in order to generate plasmid pGEX-hRap1 encoding for the wild-type hRap1 (id: Q9NYB0-1). First, the hRap1 in plasmid pLPC hRap1-FL (Addgene cat. 12542) was amplified and cloned at BamHI/XhoI restriction site of the pGEX-vector using primers hRap1-F and hRap1-R. The resulting construct was missing one entire codon for a glutamate residue within the polyglutamate stretch at positions 297 to 304. The additional codon for glutamate was inserted in this region by standard PCR using primers hRap1-Glu-F and hRap1-Glu-R.

### Protein Expression and Purification.

All the proteins were expressed in Rosetta2(DE3)pLysS cells by growing them at 37 ºC until OD600 ≈ 0.7 in LB-Miller media; then cooled down at 4 ºC for 30' before the addition of 0.7 mM of isopropyl 1-thio-β-d-galactopyranoside, followed by overnight growing at 16 ºC. Finally, cells were harvested and stored at −80 ºC.

The first steps of purification were the same for all the constructs. Briefly, cells were resuspended in lysis buffer (50 mM sodium phosphate, 0.4 M NaCl, 10% glycerol, 1 mM EDTA and 5 mM DTT at pH 7.3) supplemented with 0.1 mM PMSF. Cells were lysed by sonication, debris removed by centrifugation at 14000 rpm and the supernatant incubated overnight at 4 ºC with Glutahione Sepharose™ 4 Fast Flow (GE Healthcare). The resins were then washed with 3 volume of lysis buffer followed by an additional 3 volume wash with lysis buffer containing 1 M NaCl. Finally, the resins were equilibrated with lysis buffer and the GST-conjugated protein eluted with 25 mM reduced L-glutathione in lysis buffer. Finally, 3C-protease was added to the eluate, followed by overnight dialysis at 4 ºC against the buffer used in the next purification step.

For the TRF2 constructs, the proteins were dialyzed against buffer T (20 mM Tris-HCl, 10% glycerol, 1 mM EDTA and 1 mM DTT at pH 8) + 250 mM NaCl, applied onto a Macro-Prep High Q support (BIO-RAD) and the flow-through loaded on POROS 50 HE resin (Applied Biosystems) and eluted batch-wise with buffer T containing 400, 600 and 1000 mM NaCl. Fractions containing the purified protein were pooled, concentrated on Amicon Ultra centrifugal filters (Millipore) and dialyzed against buffer ST (20 mM Hepes, 400 mM NaCl, 40% glycerol and 1 mM EDTA at pH 7.4).

For hRap1, the protein was dialyzed against buffer T + 150 mM NaCl, loaded on Macro-Prep High Q support and eluted with buffer T + 300 mM NaCl. Pure hRap1 was concentrated with on Amicon Ultra centrifugal filters (Millipore), and dialyzed against buffer ST and stored at −80 ºC. Before use the proteins were dialyzed in buffer H (20 mM Hepes-KOH pH7.4, 4% v/v glycerol, 1 mM DTT) + 100 mM KCl (unless otherwise indicated).

All the proteins were quantified by UV using extinction coefficients at 280 nm calculated with Protparam tool (https://web.expasy.org/protparam/). For selected protein constructs the dispersity of the sample and the oligomeric state of the protein were examined by analytical sedimentation velocity using an Optima XLA (Beckman) (see Figure S1B and 1C).

*Fluorescence labeling of TRF2-Cys$^{160}$*. The protein was extensively dialyzed in buffer L (20 mM sodium phosphate pH 7.4, 400 mM NaCl, 1 mM EDTA, 10% v/v glycerol, 50 µM TCEP), concentrated with Amicon Ultra centrifugal filters to ~ 300 µL ~100 µM, followed by addition of a 10-fold molar excess of Alexa-488 maleimide and incubated at 4 °C overnight under gentle rocking. The sample was then loaded on a BioGel P6 resin equilibrated in buffer L, to separate the protein from unincorporated dye. The labeled protein was quantified spectrophotometrically to ensure stoichiometric labeling, concentrated again with Amicon Ultra centrifugal filters, dialyzed in buffer ST and quantified one last time before flash-freeze and storage at −80 °C.

**DNA Substrates.**

*DNA oligonucleotides* – The unlabeled and fluorescently labeled oligonucleotides used for the preparation of the 24 bp dsDNA substrates and oligo-$dT_n$ sequences with $21 < n < 60$ were purchased from IDT (Coralville, IA). For the unlabeled dsDNA substrates the two strands were mixed at equimolar ratio, while the FAM-labeled dsDNA substrates were annealed in the presence of 5% of the unlabeled strand. Annealing was carried out in 20 mM HEPES (pH 7.4), 100 mM NaCl, 10% (v/v) glycerol, 2 mM $MgCl_2$, by incubation in a preheated 95 °C water bath, followed by slow cooling to room temperature. Poly-dT longer than 250 nt (with a majority over 500 nt) and poly-U shorter than 250 nt were purchased from Midland Certified Reagent (Midland, TX).

*$T_2AG_3$ Cassettes* – Plasmid pSXneo 135-$T_2AG_3$ (Addgene cat. 12402) in its original Stbl3 strain was used to generate the 135-$T_2AG_3$ cassette. A sequence containing 20 *$T_2AG_3$* repeats in pUC57 was purchased from GenScript (Piscataway, NJ) and cloned into pUC19 at the EcoRI/HindIII restriction sites. During cloning a full $T_2AG_3$ repeat was lost, resulting pUC19-$(T_2AG_3)_{19}$ plasmid. The plasmid was transformed in DH5$\alpha$, purified by miniprep and sequenced several times to test the stability of the $(T_2AG_3)_{19}$ cassette in *E. Coli* and test for recombination events which may change the sequence.

All plasmids were purified following a modified Maniatis's protocol[1]. Briefly, transformed cells were grown overnight at 37 ºC, then harvested and resuspended in 50 mM glucose, 25 mM Tris-HCl (pH 8) and 10 mM EDTA. Lysis was achieved by adding 2 volumes of 1% SDS solution in 0.2 M NaOH and proteins were then precipitated with 0.35 volumes of 3 M potassium acetate in 12% acetic acid. After centrifugation, 0.6 volumes of isopropanol were added to the supernatant and the solution incubated in ice for 10' before spinning. The pellet was solubilized in TE (10 mM Tris-HCl and 1 mM EDTA, pH 8) and treated with 0.4 volumes of 10 M ammonium acetate; the precipitate was spun down, and the supernatant treated with 2 volumes of ethanol and incubated for 30' at -20 ºC before centrifugation. The pellet was resuspended in TE and the RNA contaminants were digested with RNase A for 30' at 37 ºC. NaCl (to 1.5 M final concentration) and 0.25 volumes of 30% PEG8000 were added to the solution which was centrifuged after 30' incubation in ice. The resulting pellet was solubilized in 100 mM Tris-HCL (pH 7.5), 150 mM NaCl, 12 mM EDTA and 1% SDS, then Proteinase K was added, and the solution incubated for 30' at 37 ºC. Finally, the plasmid was phenol extracted, ethanol precipitated and resuspended in TE.

pSXneo 135-$T_2AG_3$ was digested NotI-HF and SmaI while pU19-$(T_2AG_3)_{19}$ with EcoRI and HindIII to release the cassettes from the vectors. The cassettes were separated from the vector by iterative PEG8000 fractionation in TE with 600 mM NaCl. Finally, pure cassettes were ethanol precipitated, resuspended in HE and quantified. Carboxy-fluorescein labeled $^{FAM}(T_2AG_3)_{19}$ was generated by two step PCR using Phusion polymerase in GC buffer in the presence of 1 M betaine using pU19-19-$T_2AG_3$ as template. The annealing/amplification steps were carried out at 72ºC for 30 seconds with primers FAM-pUC19-842 and pUC19-929.

*DNA substrates for the magnetic tweezer* – Non-telomeric DNA was generated by Pfu PCR amplification of a 2kb region from (pGOHis4TATA) using primers with unique restriction sites (GO-His4-MluI-F, GO-StyI-R). Two DNA handles (1kb) were TAQ PCR amplified from (pSH726). One DNA handle was amplified in the presence of dUTP-biotin with primers (F-T1-DH-1kb, R-T1-DH-NheI) while the other was amplified in the presence of dUTP-digoxigenin with primers (F-T1-UH-AscI, R-T1-UH-1kb), resulting in multiple, random incorporations of label along the length of the handle DNA. PCR products were restriction digested with the appropriate restriction endonuclease and then ligated together with T4 DNA ligase following standard protocol. After each enzymatic step the DNA was purified using Qiagen PCR cleanup kit to remove contaminants. A 2:1 molar excess of DNA handles was used in the ligation reaction to drive formation of the full-length DNA substrate (~4 kbp). The full-length DNA substrate was purified by agarose gel electrophoresis (0.8% agarose in 0.5x TBE at 125 V). The gel isolated DNA was electro-eluted from the gel, purified, and concentrated using a single Qiagen spin column following manufacturer's protocol. The DNA was eluted in 10 mM Tris-HCl pH 8.0 and supplemented with glycerol and EDTA to final concentrations of 50% (v/v) and 1 mM, respectively.

DNA containing human telomeric sequence was generated by incorporating the $(T_2AG_3)_{135}$ cassette as follows. The $(T_2AG_3)_{135}$ cassette was isolated by restriction digest of pSXneo 135-$T_2AG_3$ with EcoRI and NotI. Two symmetric DNA extensions of 625 bp were PCR amplified from pUC19 using primers (pUC19-927-EcoRI / pUC19-1553r-NheI) for one extension and primers (pUC19-927-NotI / pUC19-1553r-MluI) for the other extension, following the two step PCR as described above for $^{FAM}(T_2AG_3)_{19}$. Two 1kb DNA handles were PCR amplified as described above. The full-length DNA was assembled in two ligation steps. First, the DNA handles were digested and ligated to their compatible DNA extensions (AscI:MluI and NheI:NheI). Then, the resulting ligation products were digested with EcoRI and NotI and ligated to the isolated $(T_2AG_3)_{135}$ cassette. The full-length DNA product was isolated by agarose gel electrophoresis, purified, and concentrated as described above for the non-telomeric DNA.

**Optical tweezers.**

Experiments were performed using a custom-built dual-optical-trap based on a 1064 nm solid state laser split by polarization[2]. λ–DNA was biotinylated by T7 DNA polymerase (NEB) in T7 polymerase buffer for 40' at 12 ºC in the presence of 0.6 mM dNTPs (dATP, dTTP, and dCTP) and 1 mM biotinylated-dGTP, and purified on spin column S-400 HR. A laminar flow-cell (μ-Flux, Lumicks) was used to establish DNA tethers in the absence of TRF2 by flowing biotinylated λ–DNA past trapped streptavidin-coated 2.1 μm beads (Spherotech, SVP-20-5) under constant flow. The binding of DNA to the bead was judged by the presence of an increase in the force exerted on that bead due to the increased drag of the DNA. Subsequently, the beads and DNA were moved to a DNA-free lamina and a tether was formed between the two beads by fishing for the other end of the biotinylated DNA using the second bead trapped in the second trap. Once the presence of a single λ–DNA in the dumbbell configuration was confirmed by force-extension measurements, the second trap was turned off and the DNA was held extended only by the flow force. Finally, the tether was moved into a lamina were TRF2 was present. At a constant high flow force ($\geq$ 4 pN as judged by the end-to-end distance of the DNA) the DNA remained extended. The flow force was then gradually reduced by raising the height of exit tube. At a critical flow velocity (*i.e.*, force), the end-to-end distance of the DNA as monitored by the bead-bead distance would decrease until the second bead made contact with the first bead and was trapped in the first optical trap. Movies were recorded at 32 fps and the position of the free bead over time was determined by a tracking routine in ImageJ. All experiments were performed at room temperature.

**Magnetic tweezers.**

The experiments were performed with a custom-built magnetic tweezer instrument[3]. DNA tethers were constructed to have 1 kb biotin handles on one end and 1 kb digoxygenin handles on the other. The DNA was first bound to 1 μm streptavidin-coated paramagnetic beads (MyOne, Invitrogen) in a test tube. The DNA-bead pairs were then immobilized on an anti-digoxygenin:polyethylene glycol coated surface by flowing them into the flow cell, incubating for 5 minutes and then flowing out non-bound beads. Force-extension control experiments confirmed the presence of a single DNA tethers on individual beads in buffer HK150 (20 mM Hepes-KOH pH7.4, 150 mM KCl, 2% v/v glycerol, 1 mM DTT) supplemented with 2 mg/mL BSA and 0.1% v/v Pluronic F-127 (ThermoFisher). After addition of 300 nM TRF2 on fully extended DNA held at maximum force (~ 5 pN, magnet position 0 mm), force release experiments were performed by changing the magnet position at a constant velocity (0.004 mm/sec) over 12.5 minutes until a force of ~ 0.3 pN was reached (magnet position 3 mm). Subsequently, the force was increased by gradually increasing moving the magnets back towards the sample over another 12.5 minutes. Movies of the force-extension experiments were collected, and the xyz-position of the DNA tethered beads and surface attached reference beads were tracked off-line using the NanoBLOC software[4]. Instrumental drift in the DNA tethered beads' position was accounted for by subtracting the xyz-displacement of surface attached reference beads[3]. Prior to the force-extension experiments, a force calibration measurement was collected by tracking DNA tethered beads at different magnet heights and calculating the force using the inverse pendulum model[3]. All experiments were performed at room temperature.

**Turbidity experiments.**

Equal volumes of 2X solutions of DNA and protein were mixed to yield the appropriate DNA to protein ratio, in a final buffer H2 (20 mM Hepes-KOH pH7.4, 2% v/v glycerol, 1 mM DTT) + 100 mM KCl (unless otherwise indicated). Absorbance of the solutions at 340 nm was measured using a NanoDrop 2000 (Fisher Scientific) after 1 minute of mixing. The reported error bars are from three independent readings. All the experiments were performed by maintaining constant the protein concentration and varying the DNA concentration. Unless otherwise indicated, the TRF2 concentration used in the experiments was 10 μM monomers. Because of the high affinity and the high protein concentrations used, the complexes of hRap1 with the TRF2 variants were formed by mixing the individual proteins at a 1:1 ratio and incubating for at least 10 minutes before starting the experiments. All experiments were performed at room temperature.

**Fluorescence and Differential Interference Contrast (DIC) microscopy.**

In order to obtain images and determine concentrations and changes in diffusion rates in protein-DNA condensates we employed confocal fluorescence microscopy on samples doped with either fluorescein (FAM) or Cy3 3'-labeled DNA or Alexa488 labeled full length TRF2. Confocal fluorescence measurements were performed on a Picoquant MT200 instrument (Picoquant, Germany). The microscope (Olympus IX-73, Japan) is equipped with a high numerical aperture water immersion objective (60x1.2 UPlanSApo Superapochromat, Olympus, Japan). Fluorophores were excited using a 485 nm pulsed laser (LDH P-C-485, Picoquant, Germany) with a repetition rate of 20 MHz. Excitation power was monitored before the objective with a laser photodiode and optimized to avoid photobleaching under normal measurement conditions, saturation of detectors at the maximum labeling concentration, and to get the higher signal to noise ratio, in a way that allowed us to keep a constant power for each set of measurements. Emitted photons were collected with a 60x1.2 UPlanSApo Superachromat water immersion objective (Olympus, Japan), passed through a dichroic mirror (ZT568rpc, Chroma, USA), and filtered by a 100 µm pinhole (Thorlabs, USA). Photons were separated according to polarization using a polarizer beam splitter cube (Ealing, California, USA) and further refined by a bandpass filter, either 525 nm ± 25 nm (ET525/50m, Chroma, USA) for FAM and Alexa488 fluorophores, or 642 nm ± 40 nm (ET642/80m, Chroma, USA) for Cy3, in front of the SPAD detectors (Excelitas, USA). Photons were counted and accumulated by a HydraHarp 400 TCSPC module (Picoquant, Germany) with 16 picosecond resolution.

All measurements were performed in either uncoated polymer coverslip cuvettes (Ibidi, Wisconsin, USA) or glass coverslips with 4 mm-diameter glass cylinder attached on top. Between measurements cuvette wells were soaked with hot 10% DECONEX11 Universal cleaning solution, rubbed with CleanWIPE swaps, rinsed thoroughly with distilled water, then di-destilled water and finally dried by flowing dried air. All measurements were performed at 23.0 ± 0.5 °C in a temperature-controlled room, as detected in the microscope stage.

DIC imaging of the phase-separated solutions was performed with an inverted Olympus IX81 microscope equipped with Nomarski prisms and a Hamamatsu C9100 EM-CCD camera. A drop of solution was deposited on a glass coverslip and imaged with a 10X objective as the droplets deposit by gravity onto the surface. Movies were recorded at 10 fps. Static DIC images of selected fields of view were generated in Image J by Z-average of 50 frames, and used to measure the diameter of the formed droplets. All experiments were performed at room temperature.

**Fluorescence recovery after photobleaching (FRAP) determinations.**

In order to monitor the diffusion of either TRF2 or DNA within condensates we performed fluorescence recovery after photobleaching (FRAP) experiments on selected TRF2-DNA droplets. Measurements were performed by focusing inside the droplets and photobleaching a limited area of the droplet determined by the size of the illuminated volume at the corresponding laser power.

Samples contained either fluorescently label protein or fluorescently label DNA. 30 µl samples were prepared in 1.5 ml-plastic test tubes at room temperature. For samples containing fluorescently labeled protein, unlabeled protein was first premixed with its labeled counterpart in the corresponding final buffer and salt concentration and then was mixed with DNA. For fluorescently labeled DNA, this was first premixed with its unlabeled counterpart and then mixed with protein. Within 10 min after mixing, 0.5 µl of 0.1 % v/v Tween20 and 0.5 µl of beta-mercapthoethanol were added, to reduce nonspecific sticking to cuvette walls and increase photostability[5], respectively, and samples were mixed and pipetted into plastic low binding tubes.

Otherwise indicated, fluorescence measurements were performed on samples containing either fluorescein (FAM) 3'-labeled DNA or Alexa 488 labeled full length TRF2, at a total concentration of 50 nM labeled material mixed with the concentrations of unlabeled DNA and protein indicated for each experiment.

Pre-photobleaching and after-photobleaching signals were recorded with an excitation power of 8 nW and 40 nW for FAM-labeled DNA and Alexa488-labeled TRF2, respectively. Photobleaching of focused spots inside the droplets was accomplished by 1 minute irradiation at 20 µW for FAM-labeled DNA and at 100 µW for Alexa488-labeled TRF2. Representative FRAP traces showed in *Results* correspond to individual time traces time-binned with 100 ms binwidth. All experiments were performed at room temperature.

*FRAP analysis* – To analyze FRAP traces and obtain the extent of recovery and mean diffusion times we employed the 3-dimensional FRAP formalism described by Blonk *et al*[6] solved for the condition of no scanning. According to this formalism, the relative fluorescence signal,

$$\Delta F_{rel}(t) = \frac{F(t)-F_{bkg}}{F_{pre}-F_{bkg}} \qquad \text{(eq. S1)}$$

where $F(t)$, $F_{bkg}$ and $F_{pre}$ are respectively the time dependent fluorescence intensity recorded after photobleaching, the background recorded from buffer alone and the fluorescence recorded before photobleaching, for the case of a single diffusing species is described by:

$$\Delta F_{fit}(t) = \Delta F_{max}(t)\left[1+0.5\left(1-exp\left[\frac{-2r_0^2}{\omega^2}\right]\right)^{-1} Erf\left[0,\sqrt{2}\frac{z_0}{\zeta}\right]\sum_{m=0}^{m_{max}}\frac{(-\kappa)^m}{m!\sqrt{\alpha_m}\beta_m}Erf[-Z_m,Z_m]\gamma_m\right] \qquad \text{(eq. S2a)}$$

$$Z_m = \left[\frac{2}{\left(1+m+2m\frac{t}{\tau_D}\frac{\omega^2}{\zeta^2}\right)\left(1+2m\frac{t}{\tau_D}\frac{\omega^2}{\zeta^2}\right)}\right]^{0.5}\left(1+m+2m\frac{t}{\tau_D}\frac{\omega^2}{\zeta^2}\right)\frac{z_0}{\zeta} \qquad \text{(eq. S2b)}$$

$$\alpha_m = 1+m\left(2\frac{t}{\tau_D}\frac{\omega^2}{\zeta^2}+1\right) \qquad \text{(eq. S3)}$$

$$\beta_m = 1+m\left(2\frac{t}{\tau_D}+1\right) \qquad \text{(eq. S4)}$$

$$\gamma_m = \left(1-exp\left[\frac{-2r_0^2}{\omega^2}\left(1+\frac{m}{1+2m\frac{t}{\tau_D}}\right)\right]\right) \qquad \text{(eq. S5)}$$

$$Erf[x_1,x_2] = \frac{2}{\sqrt{\pi}}\int_{x_1}^{x_2} e^{-t^2}\,dt \qquad \text{(eq. S6)}$$

where $\Delta F_{max}$ is the final value of $\Delta F(t)$ and corresponds to the mobile fraction of fluorescent macromolecule under the assumption of total photobleaching of the immobile fraction; $\kappa$ is the depth of photobleaching parameter and, in the model, is the exponential decay exponent of the fluorophore concentration in the center of the focused region, valued right before fluorescence recovery begins to be recorded; $\omega$ and $\zeta$ are the horizontal and vertical $e^{-2}$ decay distances of the excitation beam profile, $r_0$ and $z_0$ are the radius and vertical half length of the assumed detection cylinder, with $r_0$ = 322 nm and $z_0$ = 1344 nm, as estimated from the fit of a 3-dimensional Gaussian function to the intensity profile of *z*-stacked *x-y* scans of immobilized 200 nm-diameter fluorescent beads; $\omega$ was set to be equal to $r_0$, and $\zeta$ was estimated as $\zeta$ = 8 $\omega$ according to the analysis of photobleaching profiles of TRF2-$(T_2AG_3)_{19}$ droplets formed in presence of 3'FAM-$(T_2AG_3)_{19}$. Finally, $\tau_D$ is the time for getting a root mean square displacement on the *xy* plane equal to $\omega^2$ through 2-D translational diffusion with the diffusion coefficient *D* of the observed labeled molecule: $\tau_D = \omega^2/(4D)$.

Fitting of Eq. S2 to the data was performed through least-squares non-linear regression analysis on individual FRAP traces obtained from 3-6 different droplets. Mean values of parameters <*p*> and their associated standard errors *s* were computed using the variance estimates from the individual fits as statistical weights $w_i$ according to:

$$<p> = \sum_i w_i \, p_i \qquad \text{(eq. S7)}$$

$$s^2 = \frac{\frac{1}{\sum_i w_i}\sum_i w_i (p_i - <p>)^2}{\frac{\sum_i w_i^2}{(\sum_i w_i)^2} - 1} \qquad \text{(eq. S8)}$$

*Phase diagram tie-lines* – The concentration of TRF2 and DNA ligands in the two phases was determined employing the fluorescence intensity recorded in both phases under the confocal microscope, assuming a linear relationship between intensity and concentration and a constant molecular brightness (molar fluorescence) equal to that of the free labeled protein (Alexa488-TRF2) or labeled DNA (3'-FAM-DNA substrates). Concentrations were computed as

$$[X_i] = \frac{(F_i - F_{bkg})}{(F_{ref} - F_{bkg})}[X_{tot}] \qquad \text{(eq. S9)}$$

where *X* indicates either TRF2 or DNA substrate, subindex *i* indicates the light or dense phase; $F_i$, $F_{bkg}$ and $F_{ref}$ are respectively the fluorescence intensity recorded in the *i* phase, in buffer alone and in a solution of the labeled macromolecule alone -Alexa488-TRF2 or 3'FAM-DNA- at the same concentration and same laser power than in the final mixture; and $[X_{tot}]$ is the total concentration of the observed component (both labeled and unlabeled) in the final mixture.

Measurements of intensity in the dense phase was performed by focusing inside multiple droplets in 2-4 different 80 µm x 80 µm fields of view, at 2-10 µm from the bottom surface. Measurements in the light phase were performed by focusing in the free solution space between droplets or, for very crowded surfaces where was impossible to reliably focus in a spot without interference from neighboring droplets, in the supernatant pipetted to another cuvette after performing the measurements in the dense phase.

**Coarse-grained simulations.**

*Variable force constant simulations* – Coarse-grained simulations were performed using the PIMMS simulation engine[7]. Briefly, all force-dependent simulations were performed with an 80-mer "DNA" strand in an 80x80x80 box with 4000 copies of the protein bead. Extension force was applied with a distance-dependent harmonic potential with an ideal distance of 80 lattice units and a varying force constant. Practicaly speaking, individual simulations were run at under equilibrium conditions with different fixed force constants (as opposed to a *bona fide* pulling simulation), with the resulting extension vs. force constant data shown in Figure 2 and Figure S10. Protein molecules were represented as single beads, with an excluded volume of one lattice site and short- and long- range interactions that extend one or two lattice sites away from the bead in every direction. As such, the radius of gyration for the bead is 2.5 lattice units (1/2 the diameter of the volume in which interactions are 'felt), while the excluded volume is just a single lattice voxel.

At least five independent replicas were run for each possible condition, with 10 - 20 replicas for many systems. Individual replicas were run for between $2 \times 10^9$ and $9 \times 10^{10}$ Monte Carlo steps. Specifically, for force constant values in which bimodal behavior was observed, all independent replicas were run for a sufficient number of steps to ensure that multiple exchanges between the two states are observed in a single simultion. Table 8 outlines the interaction parameters used in units of $kT^{-1}$. Simulations were performed at a temperature of T=140 with with an 80:15:5 moveset ratio for crankshaft, chain translate, and chain rotate moves. Interaction parameters reported in Table 8 reflect strengths in kT, i.e. parameter file value divided by temperature (k=1).

We estimated the TRF2 solubility threshold in simulations by performing simulations in a fixed box volume at variable TRF2 concentrations and determining the concentration at which self-assembly occurs. We converted lattice units to nanometers with a conversion factor of 4.5 nm per lattice unit. This conversion places the excluded volume of TRF2 molecules at 4.5 $nm^3$, with the radius of gyration for the full-length protein of 5 nm. This conversion allows us to estimate molar concentrations by calculating number of molecules per unit volume. The solubility threshold was set at the concentration at which a monodisperse solution is no longer observed, which emerges a density of 800 molecules per 90 $nm^3$. Varying the conversion factor used here changes the value between 0.5 mM and 3 mM, such that a ~1 mM order of magnitude reflects an approximate regime. The mass concentration (~30 mg/ml) was estimated by convert 0.5 mM to mass concentration using a molecular mass of 59593.50 Da for full-length TRF2. The solubility of lysozyme was taken from the lysozyme Sigma Aldritch product information (CAS RN 12650-88-3, EC 3.2.1.17, Lysozyme from chicken egg white for Molecular Biology).

*Phase separation simulations* – Phase separation simulations were run with identical parameters, movesets, and setup as variable force constant simulations. A 10-mer (short) or 20-mer (long) DNA polymer was used and weaker vs. stronger inteaction strengths used as in the force experiments. DNA concentration varied between 400 and 2600 DNA beads (40 and 260 molecules) and between 200 and 1000 protein beads (200 and 1000 molecules). Where phase separation occurred a single protein:DNA assemly always formed and was stable for the duration of the simulation.

**Framing DNA condensation in terms of polymer physics.**

In the following, we wish to contextualize our experimental and computational observations in terms of the language of polymer physics, as to provide a more explicit connection between force experiments, chain condensation, and phase separation. The fundamental idea that the same forces driving chain compaction are responsible for phase separation stems directly from the Flory-Huggins model[8] and has been originally discussed in terms of nucleic acid condensation by Post and Zimm[9]. In the context of biomolecular condensates, recent experiments and simulations have confirmed that indeed the molecular interactions controlling the conformation of disordered proteins are the same interactions encoding for phase separation and can be quantitatively associated[7].

Different models can be constructed to describe this phenomenon within the framework of Flory-Huggins theories. The model proposed here has to be regarded as "a" solution, that does not pretend to cover all the possible case scenario, but aims to provide a physic-based argument to sustain the connection between the information infered from force spectroscopy experiments and phase-separation. Future work is required to quantitatively assess the validity of the model over other alternative descriptions.

In the spirit of the work by Post and Zimm[9], here we will describe DNA condensation by a ligand as a modulation of the intrinsic properties of the polymer representing the nucleic acid chain. In these terms, the configuration of the polymer are dictated by two- and three-body interactions. The two-body interactions account for the physical excluded volume, repulsive electrostatics (which basically can be regarded as a virtual additional excluded volume), and the attractive interactions due to the ligand binding. A simple approximation to account for the contribution of bridging events compacting a polyelectrolyte chain has been previously described by Kundragami and Muthukumar[10], where the effective excluded volume can be defined as $v = (v_0 + \beta E_{br} c_b)$. Here $v_0$ is the two-body term in absence of ligand, $\beta = 1/KT$ , $E_{br}$ is the interaction energy of the ligand and DNA monomers when forming a bridge, and $c_b$ is the concentration of the ligand. This can easily translated in the same language used by Post and Zimm by rewriting the interaction terms as: $v = v_0\left(1 + \frac{\beta E_{br} c_b}{v_0}\right) = v_0(1 - 2\chi)$ where $\chi$ is now dependent on $c_b$. However, it is easy to incorporate in $\chi$ also temperature dependences and therefore in the following we will refer to $\chi$ as a generic interaction term.

Among different models, in this context, it is convenient to formulate force experiments accordingly to the model proposed by Morrison et al.[11]. Here the extensible Hamiltonian of a polymer under force is given by

$$\beta H_F = \frac{3}{2a^2}\int_0^N ds \dot{\boldsymbol{r}}^2(s) - \beta f \int_0^N ds\, \dot{\boldsymbol{z}}(s) + \Delta_2 + \Delta_3 \qquad \text{(Eq. S10)}$$

where

$$\Delta_2 = \frac{v}{2}\int_0^N ds'\delta[\boldsymbol{r}(s) - \boldsymbol{r}(s')] \qquad \text{(Eq. S11)}$$

and

$$\Delta_3 = \frac{w}{6}\int_0^N ds \int_0^N ds' \int_0^N ds''\, \delta[\boldsymbol{r}(s) - \boldsymbol{r}(s')]\delta[\boldsymbol{r}(s') - \boldsymbol{r}(s'')] \quad \text{(Eq. S12)}$$

With $\beta = 1/KT$, $N$ being the number of monomers in the polymer chain, $z$ the direction along which the $f$ force is aligned, $\Delta_2$ and $\Delta_3$ the contribution of two- and three-body interactions ($v_0$ and $w_0$ respectively). The first term of the Hamiltonian describes the ideal chain, whereas the second term introduces the extension of the polymer along the z-axis. The third term describes two-body interactions and, in the spirit of the Post and Zimm work, these interactions will reflect not only the repulsive nature of the electrostatic in the DNA as well as the intrisic rigidity (persistence length), but also the attractive contribution induced by the binding of proteins.

A self-consistent equation for the extension of the polymer associated with $\beta H_F$ is given by:

$$\lambda^2 - 1 = \left(\frac{3}{2\pi}\right)^{3/2} \frac{v\sqrt{N}}{\lambda^3} \int_\delta^1 \frac{1-u}{\sqrt{u}} e^{-\frac{N\lambda^2\varphi^2}{2}} + \left(\frac{3}{2\pi}\right)^3 \frac{w}{\sqrt{N}\lambda^6} \int_\delta^1 du_1 \int_\delta^{1-u_1} du_2 \frac{(1-u_1-u_2)(u_1+u_2)}{u_1^{3/2}u_2^{3/2}} e^{-\frac{N\lambda^2\varphi^2}{2}} \quad \text{(Eq. S13)}$$

with $\varphi$ is a dimensionless force given by $a\beta f, \delta$ is a cut-off parameter to ensure convergence of the integrals, and $\lambda$ is the self-consistent parameter connected to the force-induced expansion of the end-to-end distance of the polymer chain. More precisely, the mean extension $\langle Z \rangle = \varphi L \lambda^2/3$ where $L$ is the contour length of the polymer.

This self-consistent equation has up to three solutions ($\lambda_C < \lambda_B < \lambda_E$) depending on the force regime (Fig. S9A). Below a critical force $\varphi_C$ the polymer chain is a collapsed globule dominated by the attractive two-body interactions and the increasing force is only moderately expanding the chain. This mirrors what observed at low force in our experiments when the nucleic acid is bound to the protein as well as the condensed state identified in simulations. Increasing the force above $\varphi_C$ leads to the appearance of two additional solutions representing a saddle point ($\lambda_B$) and an extended conformation ($\lambda_E$). The coexistence between collapsed and expanded conformations parallels what observed in both experiments and simulations at intermediate forces. Whereas in experiments this is represented by an increase in the conformational fluctuations, the underlining mechanism is clearly highlighted in simulations (Fig. 2 and S7). By further increasing the force, above a given threshold $\varphi_E$, only the most extended configuration is favored, which mirrors the final configuration state observed in simulations and experiments at high forces.

When $f$ is set to zero, the Hamiltonian simplifies to the one of a polymer chain with two- and three- body interactions in absence of force:

$$\beta H_0 = \frac{3}{2a^2}\int_0^N ds \dot{\boldsymbol{r}}^2(s) + \Delta_2 + \Delta_3 \quad \text{(Eq. S14)}$$

whose solutions can be described by the self-consistent equation[12]:

$$\alpha^2 - 1 = \frac{4}{3}\left(\frac{3}{2\pi}\right)^{3/2} \frac{v\sqrt{N}}{\alpha^3} + \left(\frac{3}{2\pi}\right)^3 \frac{B\,w}{2\alpha^6} \quad \text{(Eq. S15)}$$

Here $B$ is equal to $\frac{1}{N}\sum_{l=3}^{N}\sum_{m=2}^{l-1}\sum_{n=1}^{m-1}\frac{(l-n)}{((l-m)(m-n))^{3/2}}$.

The equivalent free energy of the single chain in solution can be written also as[12]:

$$\beta F_{chain} = \frac{3}{2}(\alpha^2 - 2\ln\alpha) + \left(\frac{3}{2\pi}\right)^{3/2} \frac{v\sqrt{N}}{\alpha^3} + \left(\frac{3}{2\pi}\right)^3 \frac{B\,w}{2\alpha^6} \quad \text{(Eq. S16)}$$

When mixing multiple chains, the free energy of the solution can be written in terms of the number of solvent and polymer molecules $n_1$ and $n_2$ and their corresponding volume fractions $v_1$ and $v_2$:

$$\beta F_{mix} = n_1 \ln v_1 + n_2 \ln v_2 + \chi n_1 v_2 \quad \text{(Eq. S16)}$$

In the work of Post and Zimm[9] the two free energies are then combined as:

$$\beta F_{tot} = \beta F_{mix} + n_2\,\beta F_{chain} = n_1 \ln v_1 + n_2 \ln v_2 + \chi n_1 v_2 + n_2 \left(\frac{3}{2}(\alpha^2 - 2\ln\alpha) + \left(\frac{3}{2\pi}\right)^{3/2} \frac{v_0(1-2\chi)\sqrt{N}}{\alpha^3} + \left(\frac{3}{2\pi}\right)^{3} \frac{B\,w}{2\alpha^6}\right) \quad \text{(Eq. S17)}$$

The phase separation is obtained by equating the chemical potential in the light and dense phase (Fig. S9B), with the chemical potentials of the two phases being:

$$\beta(\mu_2 - \mu_{20})' = \beta(\mu_2 - \mu_{20})'' \quad \text{(Eq. S18a)}$$

and

$$\beta(\mu_1 - \mu_{10})' = \beta(\mu_1 - \mu_{10})'' \quad \text{(Eq. S19a)}$$

Here, the sub-indexes 1 and 2 refers to the solvent and polymer, $\mu_{20}$ and $\mu_{10}$ are the chemical potentials of reference of the pure species, and the upper indexes refer to the values in the two phases.

The balance of chemical potential enables reconstructing a phase-diagram of the polymer concentration as function of $\chi$. It is important to notice that this representation (as in the original Post and Zimm work[9]) does not account for the ligand concentration dependence of $\chi$, nor for the volume fraction occupied by the ligand in solution nor for the different modes of binding. This does not mean that the general hypothesis is wrong, as shown by the simulations in the present work, where the same set of interactions dictates both the collapse of the single chain (and its response to force) and the phase separation of multiple chains. A quantitative description of the phenomenon requires to expand the model accounting for these additional elements, which can be derived in the context of a three-component Flory-Huggins scheme. Advancement in the single-molecule spectroscopy of nucleic acid condensation has led to develop new models that accounts for the effect of surface tension and capillary forces (which can bring together distant parts of the nucleic acid[13]) as well as the contribution of pre-wetting near a surface[14].

**Supplementary references.**


1 Sambrook, J., Fritsch, E. F. & Maniatis, T. *Molecular cloning: a laboratory manual*. (Cold Spring Harbor Laboratory Press, 1989).

2 Moffitt, J. R., Chemla, Y. R., Smith, S. B. & Bustamante, C. Recent advances in optical tweezers. *Annu Rev Biochem* **77**, 205-228, doi:10.1146/annurev.biochem.77.043007.090225 (2008).

3 Galburt, E. A. in *Handbook of Imaging in Biological Mechanics* Ch. 38, 481-449 (CRC Press, 2014).

4 Cnossen, J. P., Dulin, D. & Dekker, N. H. An optimized software framework for real-time, high-throughput tracking of spherical beads. *Rev Sci Instrum* **85**, 103712, doi:10.1063/1.4898178 (2014).

5 Schuler, B., Müller-Späth, S., Soranno, A. & Nettels, D. in *Intrinsically disordered protein analysis* 21-45 (Springer, 2012).

6 BLONK, J. C. G., DON, A., Van AALST, H. & BIRMINGHAM, J. J. Fluorescence photobleaching recovery in the confocal scanning light microscope. *Journal of Microscopy* **169**, 363-374, doi:https://doi.org/10.1111/j.1365-2818.1993.tb03312.x (1993).

7 Martin, E. W. *et al.* Valence and patterning of aromatic residues determine the phase behavior of prion-like domains. *Science* **367**, 694, doi:10.1126/science.aaw8653 (2020).

8 Rubinstein, M., & Colby, R. H. . *Polymer physics*. (Oxford: Oxford University Press, 2003).

9 Post, C. B. & Zimm, B. H. Theory of DNA condensation: collapse versus aggregation. *Biopolymers* **21**, 2123-2137, doi:10.1002/bip.360211104 (1982).

10 Kundagrami, A. & Muthukumar, M. Theory of competitive counterion adsorption on flexible polyelectrolytes: divalent salts. *J Chem Phys* **128**, 244901, doi:10.1063/1.2940199 (2008).

11 Morrison, G., Hyeon, C., Toan, N. M., Ha, B.-Y. & Thirumalai, D. Stretching Homopolymers. *Macromolecules* **40**, 7343-7353, doi:10.1021/ma071117b (2007).

12 Huihui, J., Firman, T. & Ghosh, K. Modulating charge patterning and ionic strength as a strategy to induce

conformational changes in intrinsically disordered proteins. *J Chem Phys* **149**, 085101, doi:10.1063/1.5037727 (2018).

13 Quail, T. *et al.* Force generation by protein–DNA co-condensation. *Nature Physics* **17**, 1007-1012, doi:10.1038/s41567-021-01285-1 (2021).

14 Zhao, X., Bartolucci, G., Honigmann, A., Jülicher, F. & Weber, C. A. Thermodynamics of wetting, prewetting and surface phase transitions with surface binding. *New Journal of Physics* **23**, 123003, doi:10.1088/1367-2630/ac320b (2021).

**Table 1**. List of primers used for cloning and site-specific mutagenesis.

| Primer | Sequence |
|---|---|
| hTERF2-F | GGCTGCAGGAATTCGGAC |
| hTERF2-R | GGAAACTCGAGCCTGTTTCAGTTCATGCCAAG |
| h-TERF2-frame-F | GGAATTCGGCACAGGGACG |
| h-TERF2-frame-R | CGTCCCTGTGCCGAATTCC |
| hTERF2-Ala$^{477}$-F | CAAGTTCAGGCAGCTCCAGATGAAGACAG |
| hTERF2-Ala$^{477}$-R | CTGTCTTCATCTGGAGCTGCCTGAACTTG |
| hTERF2Δ42-F | TTATTGAATTCATGGCGGGAGGAGGCGG |
| hTERF2Δ86-F | TTATTGAATTCGAGGCACGGCTGGAAGAGGCAGTCAAT |
| pGEX-F | CCAAAATCGGATCTGGAAGTTCTGTTC |
| hTERF2-S294Stop-F | CCGCTGCCTCATGAACAGGGAAGGAAG |
| hTERF2-S294Stop-R | CTTCCTTCCCTGTTCATGAGGCAGCGG |
| hTERF2-A2-F | CGAATTGGATCCGCCGC |
| hTERF2-G86-R | GCTTACTCGAGTTAACCACAACCACGTTC |
| hTERF2-C148S-F | GGTTATGCAGTCTCTGTCGCGGATT |
| hTERF2-C148S-R | AATCCGCGACAGAGACTGCATAACC |
| hTERF2-C207S-F | GCTGCTGTCATTATTTCTATCAAAAACAAAG |
| hTERF2-C207S-R | CTTTGTTTTTGATAGAAATAATGACAGCAGC |
| hRap1-F | AAGTAGGATCCATGTCATTTACATTCACCAAAAGCG |
| hRap1-R | TAGAAGTCGACCAGAGATGCTCGGCAATTTAAGAAG |
| hRap1-Glu-F | TGAGAGGATCCATGGCGGAGGCGATGGATT |
| hRap1-Glu-R | TATCTCTCGAGTTATTTCTTTCGAAATTCAATCCTCCGAG |
| FAM-pUC19-842 | CAGTCACGACGTTGTAAAACGACG |
| pUC19-929 | GGAAACAGCTATGACCATGATTACG |
| pUC19-927-EcoRI | AGTCAGGAATTCCGAAACCCGACAGGACTATAAAGATACCAG |
| pUC19-927-NotI | CAAGTCGCGGCCGCCGAAACCCGACAGGACTATAAAGATACCAG |
| pUC19-1553r-MIul | AAAGGACGCGTGTGAAGATCCTTTTTGATAATCTCATGACCAAAATC |
| pUC19-1553r-NheI | AAAGGGCTAGCGTGAAGATCCTTTTTGATAATCTCATGACCAAAATC |
| GO-His4-MluI-F | GAGAGTACGCGTTCCTTGTGATGCTCGTCAGGG |
| GO-StyI-R | GAGAGACCTAGGGGTGAGCAAGAACAGGAAGG |
| F-T1-DH-1kb | GGATCATGTAACTCGCCTTGATCGTTGGG |
| R-T1-DH-NheI | GACACAGCTAGCATACCTGTCCGCCTTTCTCC |
| F-T1-UH-AscI | GAGAGAGGCGCGCCATGTAACTCGCCTTGATCG |
| R-T1-UH-1kb | CCGGATACCTGTCCGCCTTTCTCCCTTCG |
| $(T_2AG_3)_2$ | TCAGTCTTAGGGTTAGGGTTGAGC |
| mixed | TCGATACACTCAGCTCAGGAGTTC |

**Table 2**. FRAP of TRF2 in a $(T_2AG_3)_2$ and TRF2 mixture: 10 μM (mon) unlabeled TRF2 mixed with 50 nM A488-TRF2 and indicated concentration of dsDNA.

| $(T_2AG_3)_2$ and TRF2 – labeled TRF2 | | | | |
|---|---|---|---|---|
| **[dsDNA] (μM bp)** | **recovery** | ***mobile fraction*** | **K** | **τ (s)** |
| 0 | 0.02 ± 0.01 | 0.03 ± 0.01 | 3 ± 1 | 131 ± 70 |
| 15 | 0.012 ± 0.004 | 0.018 ± 0.004 | 2.7 ± 0.2 | 168 ± 60 |
| 48 | 0.06 ± 0.02 | 0.07 ± 0.02 | 5.1 ± 0.2 | 51 ± 10 |
| 120 | 0.49 ± 0.08 | 0.49 ± 0.08 | 5.89 ± 0.02 | 120 ± 40 |
| 300 | 0.7 ± 0.1 | 0.8 ± 0.1 | 5.0 ± 0.3 | 170 ± 10 |
| 600 | 0.04 ± 0.02 | 0.08 ± 0.03 | 1.51 ± 0.02 | 370 ± 200 |

**Table 3**. FRAP of labeled DNA in a $(T_2AG_3)_2$ and TRF2 mixture: 50 nM labeled DNA mixed with indicated dsDNA concentration and 10 μM (mon) unlabeled TRF2.

| $(T_2AG_3)_2$ and TRF2 – labeled DNA | | | | |
|---|---|---|---|---|
| **[dsDNA] (uM bp)** | **recovery** | ***mobile fraction*** | **K** | **tau (s)** |
| 0 | 0.18 ± 0.04 | 0.19 ± 0.04 | 5.6 ± 0.2 | 15 ± 3 |
| 15 | 0.71 ± 0.08 | 0.72 ± 0.07 | 5.90 ± 0.03 | 91 ± 25 |
| 48 | 0.97 ± 0.03 | 0.97 ± 0.03 | 5.89 ± 0.04 | 49 ± 10 |
| 120 | 0.93 ± 0.08 | 0.9 ± 0.07 | 5.7 ± 0.1 | 24 ± 9 |
| 300 | 0.96 ± 0.02 | 0.96 ± 0.02 | 5.5 ± 0.3 | 15 ± 3 |
| 600 | 0.32 ± 0.3 | 0.63 ± 0.20 | 0.70 ± 0.20 | 11 ± 6 |

**Table 4**. FRAP of TRF2 in a $(T_2AG_3)_{19}$ and TRF2 mixture: 5 μM (mon) unlabeled TRF2 mixed with 50 nM A488-TRF2 and indicated concentration of dsDNA.

| $(T_2AG_3)_{19}$ and TRF2 – labeled TRF2 | | | | |
|---|---|---|---|---|
| **[dsDNA] (μM bp)** | **recovery** | ***mobile fraction*** | **K** | **tau (s)** |
| 100 | 0.07 ± 0.03 | 0.16 ± 0.03 | 1.1 ± 0.3 | 120 ± 140 |
| 250 | 0.16 ± 0.10 | 0.26 ± 0.10 | 1.7 ± 0.2 | 276 ± 60 |

**Table 5**. FRAP of TRF2 in a $(T_2AG_3)_{19}$ and TRF2 mixture in presence of hRap1: 5 μM (mon) unlabeled TRF2 mixed with 50 nM A488-TRF2, 5 μM hRAP1, and indicated concentration of dsDNA.

| $(T_2AG_3)_{19}$ and TRF2 + 5 μM hRap1 – labeled TRF2 | | | | |
|---|---|---|---|---|
| **[dsDNA] (μM bp)** | **recovery** | ***mobile fraction*** | **K** | **tau (s)** |
| 44 | 0.42 ± 0.10 | 0.48 ± 0.10 | 3.6 ± 0.9 | 330 ± 50 |
| 100 | 0.32 ± 0.04 | 0.45 ± 0.04 | 2.0 ± 0.2 | 176 ± 30 |
| 250 | 0.31 ± 0.30 | 0.38 ± 0.30 | 3.1 ± 0.3 | 312 ± 60 |

**Table 6**. FRAP of labeled DNA in a $(T_2AG_3)_{19}$ and TRF2 mixture: 50 nM labeled FAM-DNA mixed with indicated dsDNA concentration, 5 μM (mon) unlabeled TRF2.

| $(T_2AG_3)_{19}$ and TRF2 – labeled DNA | | | | |
|---|---|---|---|---|
| **[dsDNA]unl (μM bp)** | **recovery** | ***mobile fraction*** | **K** | **tau (s)** |
| 44 | 0.02 ± 0.02 | 0.04 ± 0.02 | 1.6 ± 0.3 | 290 ± 40 |
| 100 | 0.06 ± 0.01 | 0.08 ± 0.01 | 3.7 ± 0.6 | 570 ± 80 |
| 250 | 0.4 ± 0.1 | 0.44 ± 0.10 | 4.9 ± 0.8 | 1400 ± 200 |

**Table 7**. FRAP of labeled DNA in a $(T_2AG_3)_{19}$ and TRF2 mixture: 50 nm labeled FAM-DNA mixed with indicated dsDNA concentration, 5 μM (mon) unlabeled TRF2, and 5 μM hRap1.

| $(T_2AG_3)_{19}$ and TRF2 + 5 μM hRap1 – labeled DNA | | | | |
|---|---|---|---|---|
| **[dsDNA]unl (μM bp)** | **recovery** | ***mobile fraction*** | **K** | **tau (s)** |
| 44 | 0.247 ± 0.005 | 0.280 ± 0.004 | 4.72 ± 0.03 | 480 ± 10 |
| 100 | 0.58 ± 0.01 | 0.60 ± 0.01 | 5.58 ± 0.04 | 750 ± 20 |
| 250 | 0.717 ± 0.006 | 0.752 ± 0.005 | 4.61 ± 0.07 | 405 ± 10 |

**Table 8.** Interaction strengths used for modeling (in units of kT).

| Coarse-grained simulation parameters | | | |
|---|---|---|---|
| **Bead 1** | **Bead 2** | ***Standard Strength (short/long)*** | **Weakened Strength (short/long)** |
| PROTEIN | PROTEIN | 0.143 / 0.07 | 0.136 / 0.064 |
| PROTEIN | DNA | 0.214 / 0.071 | 0.207 / 0.064 |
| PROTEIN | SOLVENT | 0 / 0 | 0 / 0 |
| DNA | PROTEIN | 0.214 / 0.071 | 0.207 / 0.064 |
| DNA | DNA | 0 / 0 | 0 / 0 |
| DNA | SOLVENT | 0 / 0 | 0 / 0 |

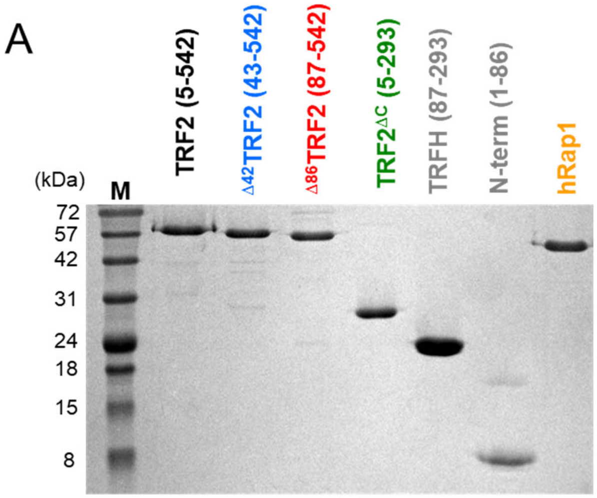


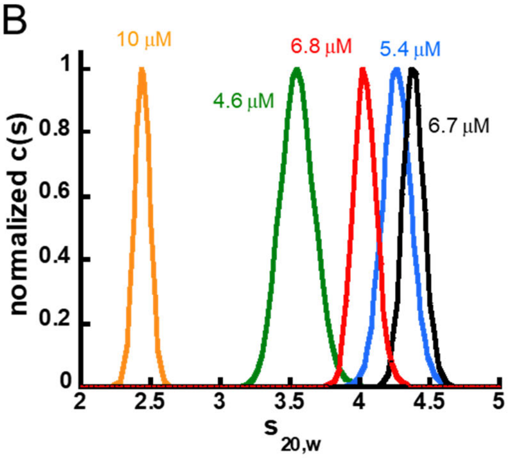


C

| | $MW_{predicted}$ (kDa) | $Mw_{calculated}$ (kDa) |
|---|---|---|
| TERF2 (5-542) | 60.1 | 112.2 |
| Δ42TERF2 (43-542) | 56.5 | 101.8 |
| Δ86TERF2 (87-542) | 52.3 | 91.8 |
| TERFΔC (5-293) | 32.8 | 64.1 |
| hRap1 | 44.7 | 44.5 |

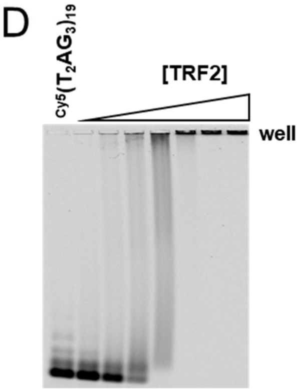


**Figure S1. A.** SDS-PAGE of representative protein constructs used in this study, stained with Coomassie Blu. **B**. Distribution of sedimentation coefficients from sedimentation velocity experiments of different protein constructs, at the indicated concentrations (in monomer). **C.** Molecular weights calculated from the sedimentation coefficients, compared to the ones predicted from the amino acid sequence. **D.** Binding of TRF2 to a Cy5 labeled DNA fragment containing 19 $T_2AG_3$ repeats, as monitored by agarose gel electrophoresis. The position of the well is indicated.

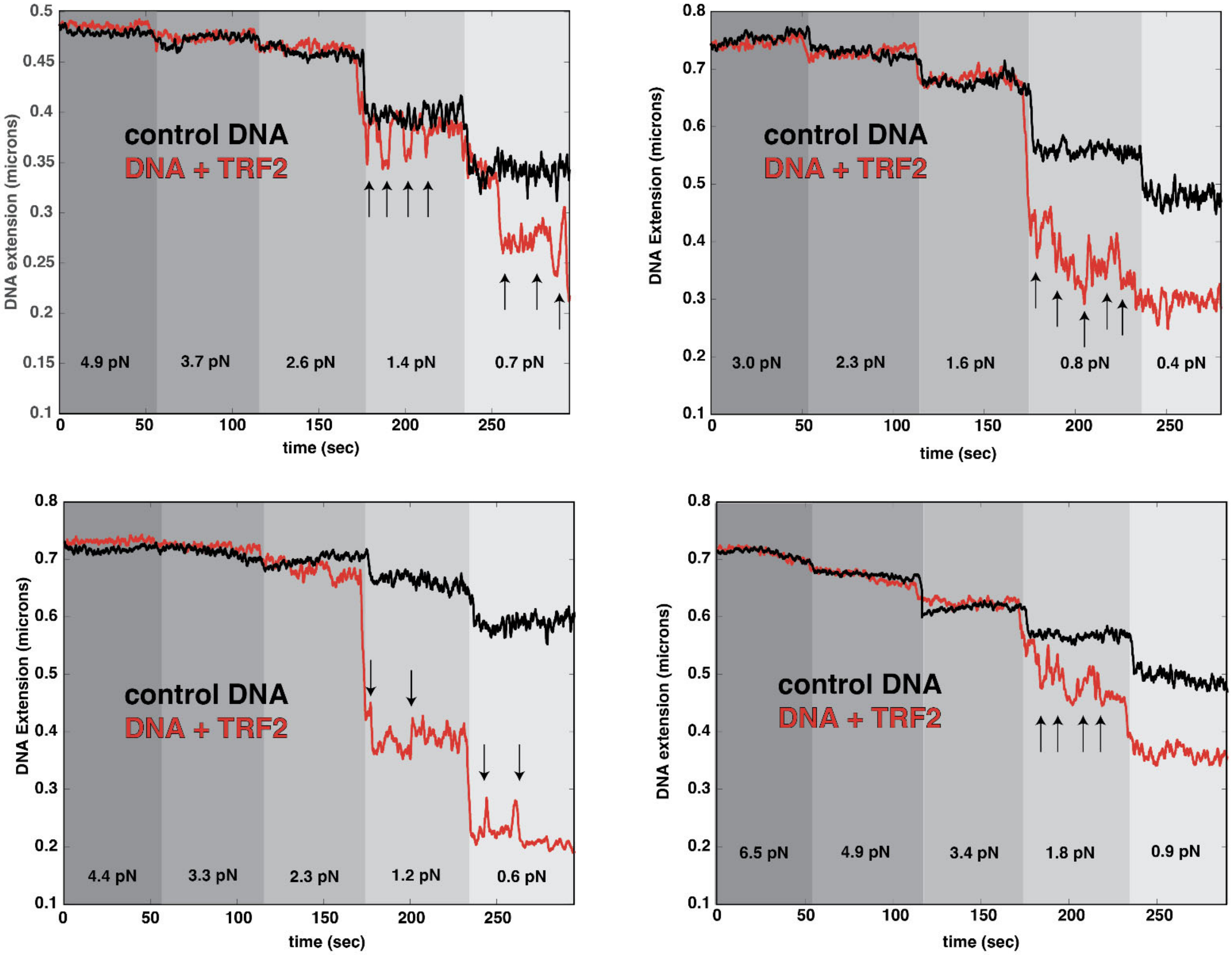


**Figure S2.** Examples of tethers where rapid transitions between metastable lengths are observed (i.e., hopping). The different shadings indicate different forces probed. These vary from tether to tether due to variation in the size of the magnetic beads. The black traces are controls where the DNA tether alone was taken through these force regimes. As expected, the length of the tether reduces as force is lowered. The red traces are experiments where TRF2 is present. Features of these traces consistent with a hopping behavior are indicated with arrows.

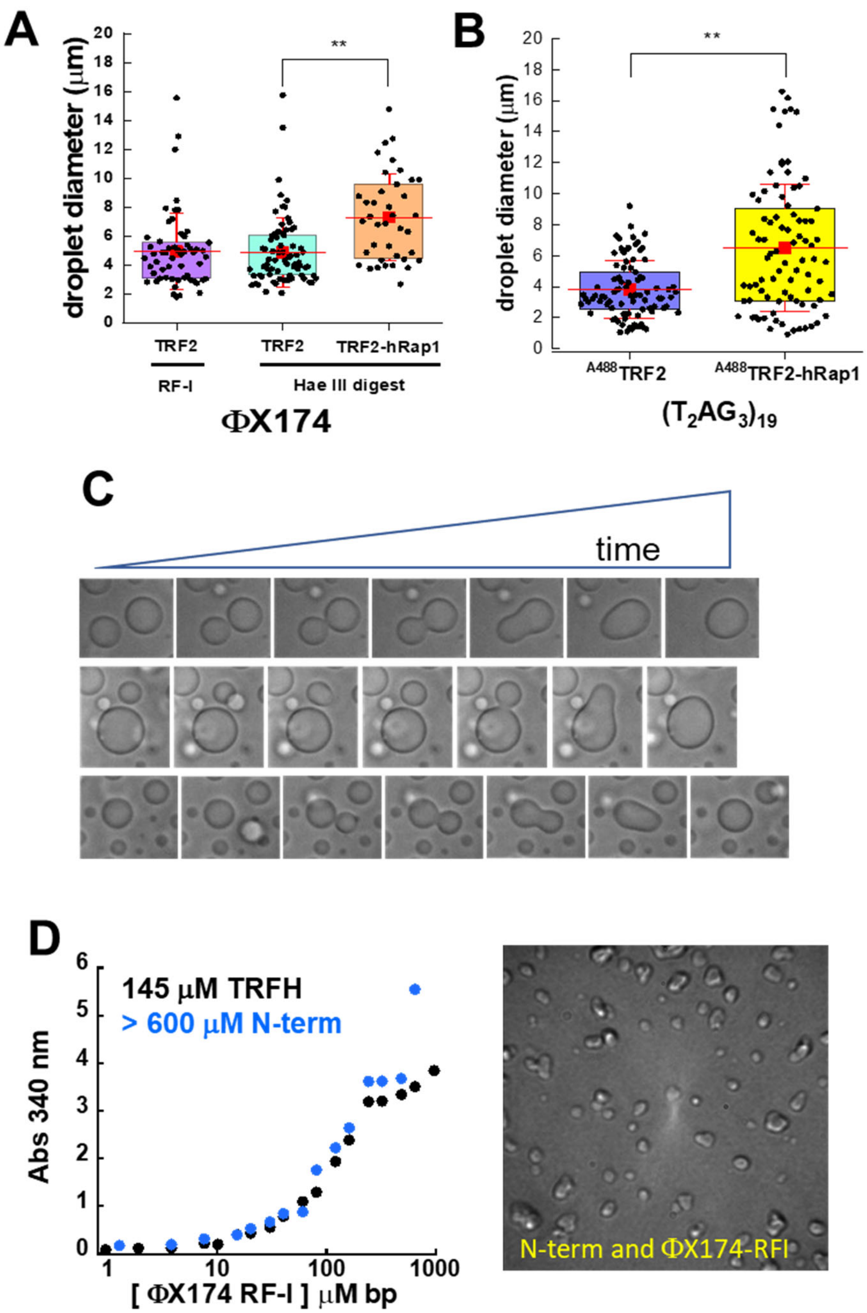


**Figure S3. A.** Diameter of TRF2-DNA droplets as observed by DIC, before the solution on the cover slip dried out. The diameter of the droplets formed in the presence of hRap1 is shown for comparison. **B.** Diameter of TRF2-DNA droplets in the absence and presence of hRap1, as monitored by fluorescence confocal microscopy via the fluorescence of Alexa-488 labeled TRF2. **C.** Example of TRF2-DNA droplets fusing on the surface. **D.** Mixing at high concentrations either the TRFH (black) or the N-terminal tail (blue) with ϕX174-RFI gives rise to an increase in turbidity, but corresponding objects formed in solutions are small amorphous droplets, substantially different from the round droplets observed with the full-length protein or other truncation variants.

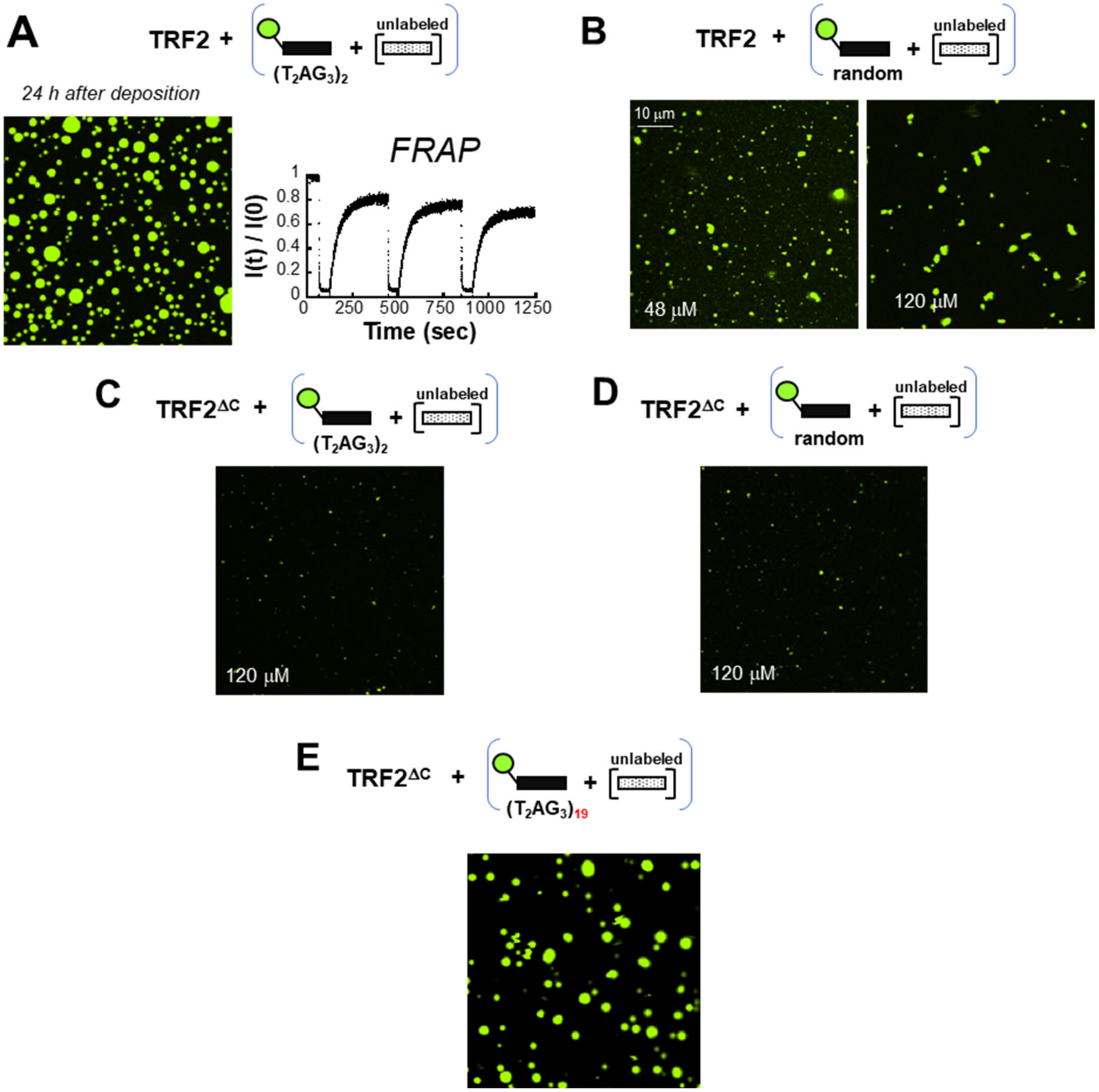


**Figure S4. A.** Droplets of TRF2 with a fluorescein-labeled 24 bp $(T_2AG_3)_2$ dsDNA after 24 h deposition on the surface maintain their liquid character, as monitored by FRAP. **B.** TRF2 does not efficiently form droplets with the fluorescein labeled 24 bp random DNA. **C, D.** $TRF2^{\Delta C}$ does not form droplets with either the $(T_2AG_3)_2$ or random 24 bp dsDNA at 120 µM. Few fluorescence puncta are observed, which are consistent with a significant shift of phase boundaries toward higher nucleic acid concentrations and a strong reduction of phase separation propensity. **E.** $TRF2^{\Delta C}$ forms droplets with the $(T_2AG_3)_{19}$ dsDNA fragment.

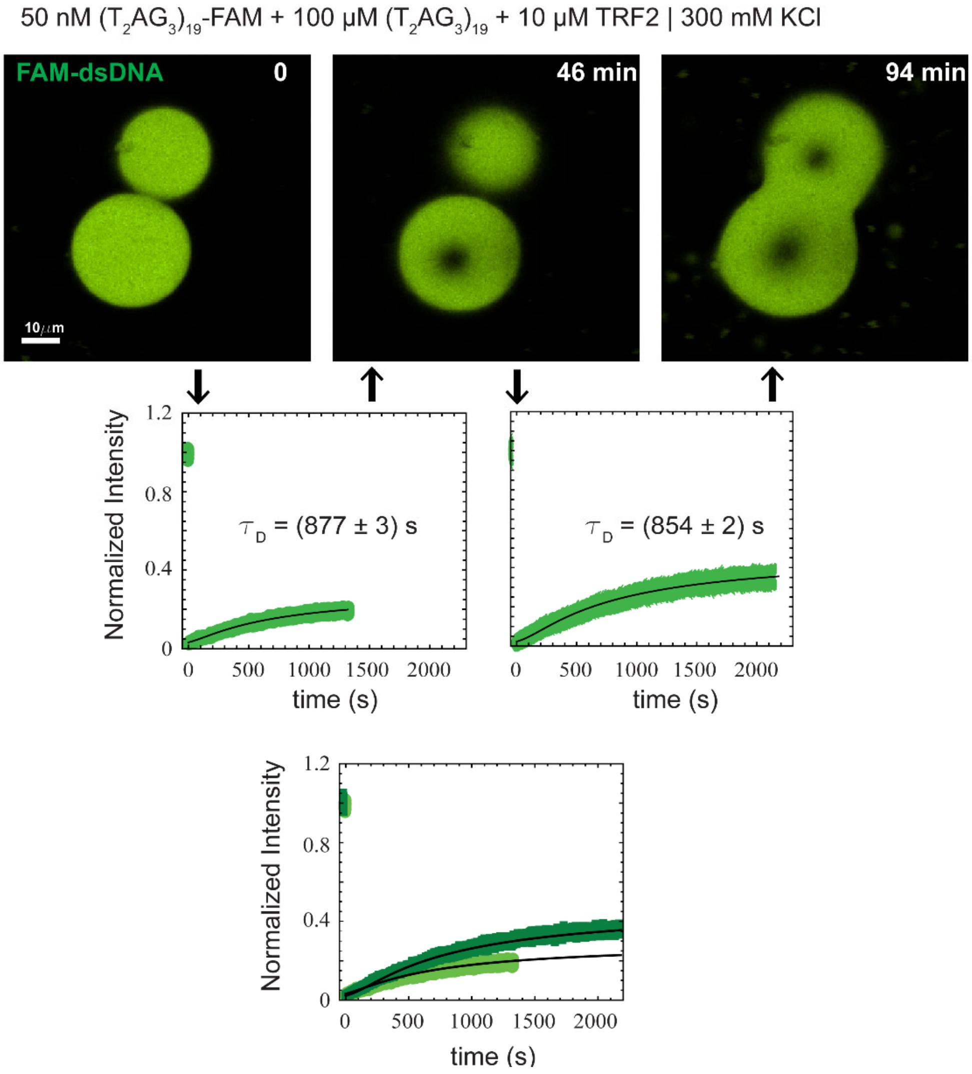


**Figure S5.** Fusion between droplets formed by 100 μM $(T_2AG_3)_{19}$ and 10 μM TRF2 at 300 mM KCl. A partial slow recovery after photobleaching is observed in each droplet with a characteristic time of approximately 800-900 seconds. Though fusion time is likely to be slowed down by interaction with the surface, this observation suggests that despite the slow and partial recovery of each droplet, the two can still fuse together.

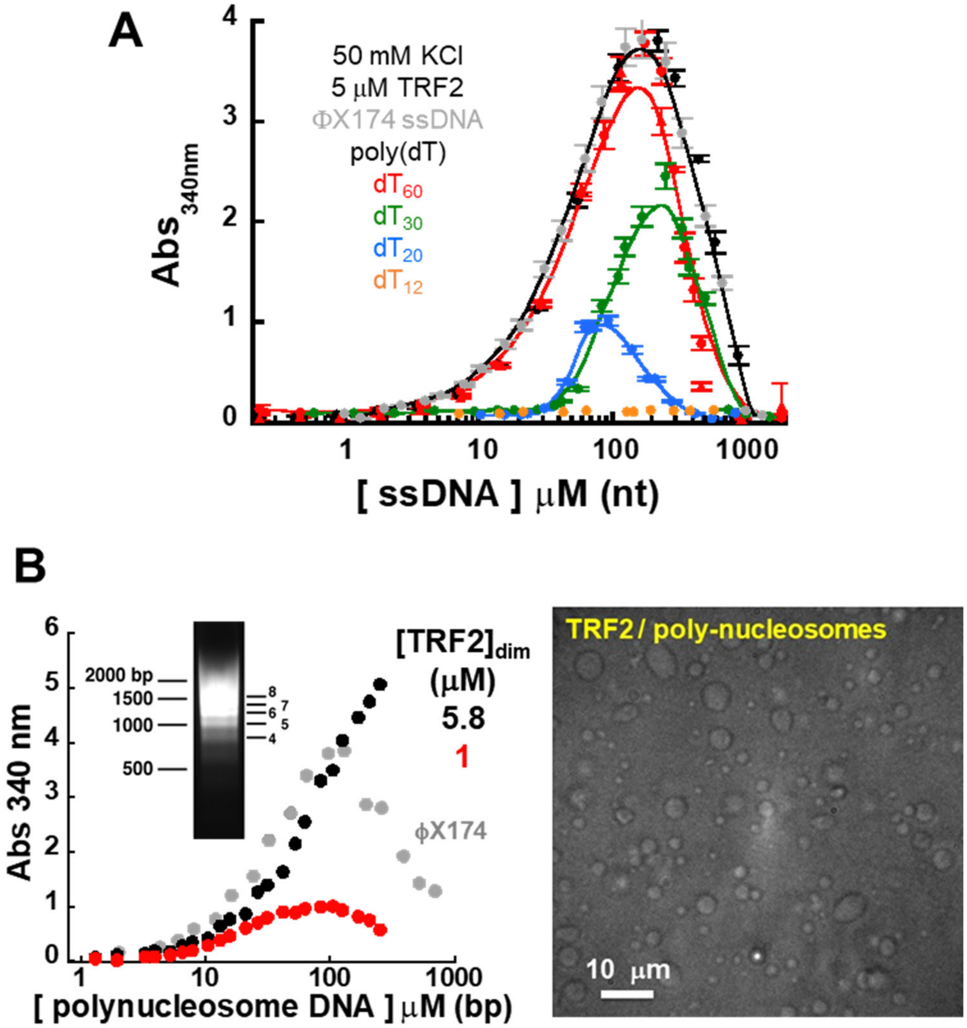


**Figure S6. A.** Absorbance scattering of TRF2 (5 μM dimer) and ssDNA of different lengths. **B.** Absorbance scattering and DIC imaging indicate that formation of poly nucleosomes does not suppress the ability of TRF2 to phase separate with DNA.

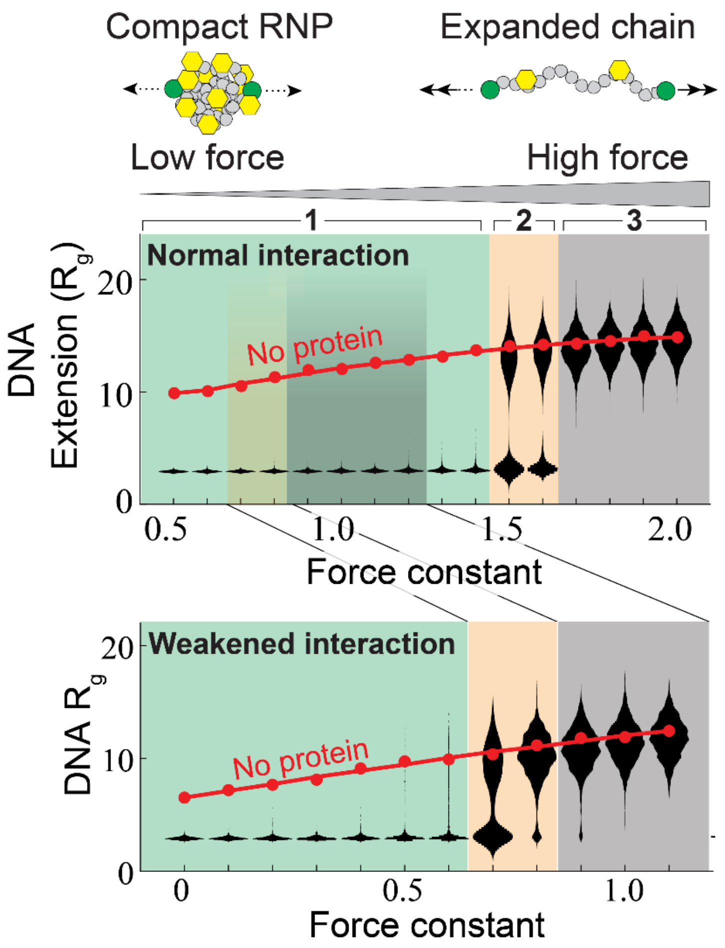


**Figure S7. Comparison of DNA extension as a function of force with violin plots.** Simulations of DNA expansion under constant force in absence of TRF2 (red dots) and in presence of the TRF2 (black data) with normal (top) and weakened (bottom) molecular interactions. In absence of TRF2 the DNA is expanded and adopts even more expanded configurations with increasing force. With a force constant of zero, the presence of TRF2 causes DNA compaction. As the force constant is increased, the DNA compaction transitions through a regime where the DNA is fully compact (region 1, top), a regime in which a bi-modal population of compact and extended molecules is realized by individual molecules oscillating between these two states (region 2, top), and eventually a regime in which the chain is fully extended (region 3, top). The transition between these three regions depends on the interaction strengths, which could experimentally be altered by adding hRap1 or by changing salt concentration. At high forces and at the transition large fluctuations occur and coexistence of the collapse and expanded state can be visualized, reflecting the large uncertainty observed experimentally in the transition. For convenience we refer to the molecules in our simulations as “DNA” and “TRF2”, but these are highly simplified models such that this language is provided as a rhetorical tool, as opposed to a specific description.

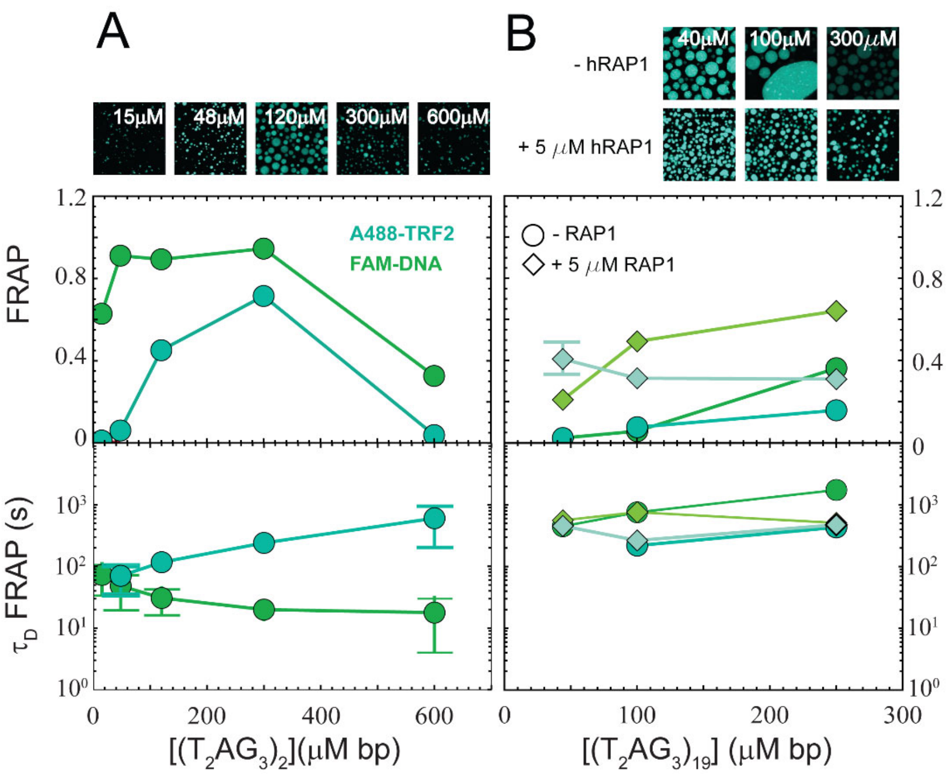


**Figure S8. Fluorescence Recovery After Photo-Bleaching amplitudes and diffusion times. A.** TRF2 phase separation with $(T_2AG_3)_2$ at increasing concentrations of specific DNA. Fluorescence Recovery After Photobleaching and diffusion time associated with the protein and DNA. **B.** TRF2 phase separation with $(T_2AG_3)_{19}$ at increasing concentrations of specific DNA. Error bars are standard deviations from FRAP experiments on different droplets at each condition.

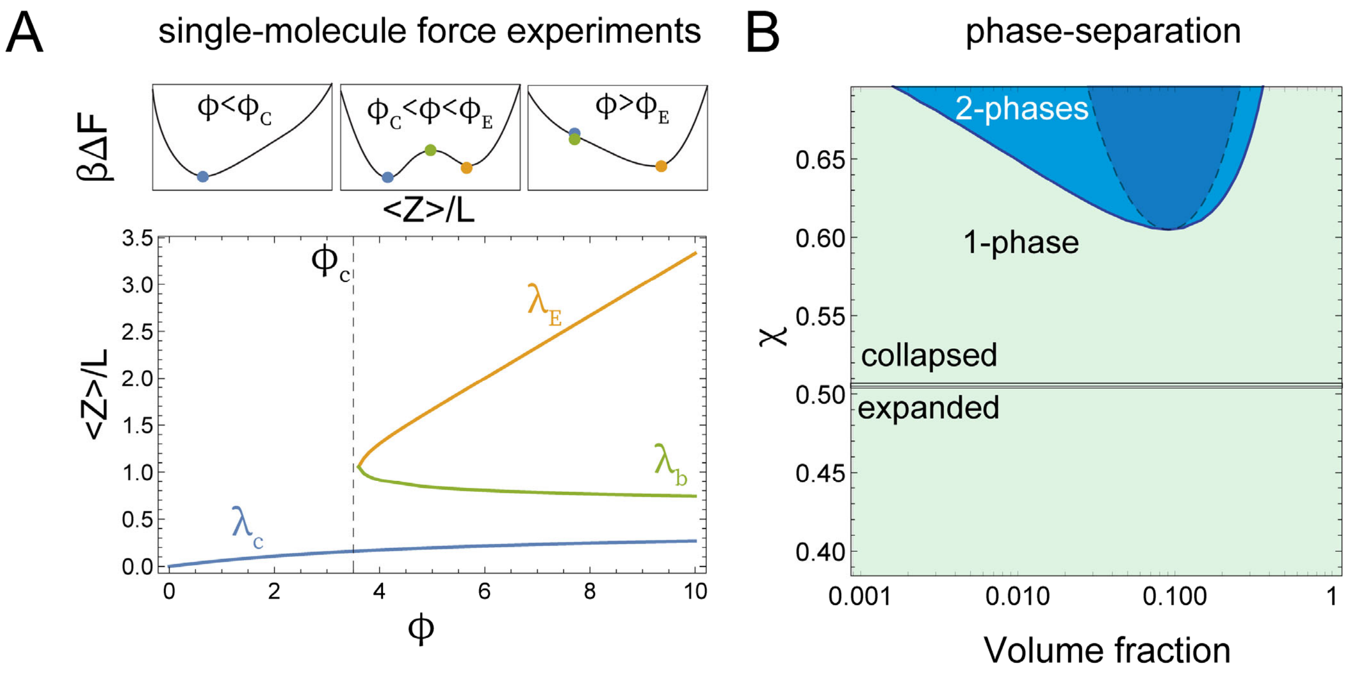


**Figure S9. Polymer theory models. A.** Single-molecule force experiments and simulations can be interpreted in terms of a single-chain force extension model[11] that accounts for two- and three-body interactions in the chain (see Eq. S10-S13). The mean extension of the end-to-end distance along the direction of the pulling force <Z>/L as function of the normalized force $\varphi$ results in three distinct solutions $\lambda_c$, $\lambda_b$, $\lambda_E$ representing a collapsed configuration, a saddle point, and an extended configuration, respectively. Increasing the force above specific threshold values ($\varphi_C$ and $\varphi_E$) favors one configuration among the others (upper panels). **B**. The same interactions that control the degree of compaction as a function of the force are responsible for the phase separation of multiple polymer chains in solution according to equations S17. A critical value of $\chi$ determines whether single chains adopts extended (good solvent) or collapsed configurations (poor solvent). In poor solvent, multiple chains within certain boundaries of concentrations can partition in two distinct phases. The light blue area represents the binodal region, whereas the dark blue represent the spinodal region. All calculations performed assuming a chain constituted by 100 monomers.
